# Pathological mutation in SMN impairs modulation of GAR1 phase separation linking condensate dysfunction to Spinal Muscular Atrophy

**DOI:** 10.1101/2025.01.30.635772

**Authors:** Sara S. Félix, Philip O’Toole, Nuno A. S. Oliveira, Brígida R. Pinho, Thomas S. Blacker, Fábio Fernandes, Jorge M. A. Oliveira, Javier Oroz, Douglas V. Laurents, Eurico J. Cabrita

## Abstract

Pathological mutations in liquid-liquid phase separation (LLPS) scaffold proteins have been linked to biomolecular condensate dysfunction in neurodegenerative diseases, while the possible impact of client protein mutations remains unclear. In spinal muscular atrophy (SMA), the disease-associated E134K mutation in the Tudor domain (TD) of the survival motor neuron (SMN) protein disrupts its interaction with GAR1, an RGG-rich protein within the H/ACA small nucleolar ribonucleoprotein complex. The consequences of this impaired interaction have not been elucidated. Here, we identified GAR1 as an LLPS scaffold protein that is phase separated in nuclear compartments and forms gel-like condensates via complex coacervation with RNA *in vitro*. In cells, we reveal that SMN co-localizes with GAR1 in Cajal bodies and modulates its dynamics. Using confocal microscopy and NMR spectroscopy, we further show that SMN TD is a client protein that regulates the architecture and dynamics of GAR1 condensates through competitive RNA interactions, implicating GAR1 phase separation in RNA accumulation and release processes regulated by SMN. Notably, the SMA-associated E134K variant of SMN exhibits a reduced affinity for GAR1, impairing the modulation of GAR1 condensates and displacement RNA. Our findings suggest a mechanistic link between phase separation dysregulation and SMA, driven by disrupted scaffold-client interactions, that highlights the therapeutic potential of targeting SMN-dependent condensate regulation in SMA.

## Introduction

Spinal muscular atrophy (SMA) is an autosomal recessive disorder characterized by the progressive degeneration of motor neurons, which are specialized nerve cells responsible for controlling both voluntary and involuntary muscle movements ^1^. Although relatively rare, SMA represents the leading hereditary cause of infant mortality ^2^. Clinically, SMA is categorized into five main subtypes, from 0 to IV, according to the severity of symptoms and disease onset ^3^. Approximately 95% of SMA cases are attributed to homozygous deletions or point mutations in the *SMN1* gene ^4^. This gene encodes the survival motor neuron (SMN) protein, which is essential for the development, maintenance, and function of motor neurons ^5^. SMN is a 38 kDa multifunctional protein that harbors several functional domains, including the N-terminal lysine-rich region, the central conserved Tudor domain (TD), the C-terminal proline-rich region, and the tyrosine-glycine (YG)-box ^6^. The TD is a conserved motif that mediates interactions between SMN and RGG/RG domains of target proteins, motifs that are frequently present in phase-separating proteins and play a significant role in liquid-liquid phase separation (LLPS) ^7–13^. The involvement of protein LLPS in SMA pathophysiology remains understudied. However, an intriguing pattern emerges, as mutations linked to the most severe forms of the disease tend to cluster within the TD of SMN, compromising the interaction between SMN and RGG-rich proteins ^12,14,15^. While this molecular disruption is considered a key feature in SMA and suggests a potential link between SMA and LLPS, the molecular mechanisms and phenotypic consequences remain to be fully elucidated.

The glycine-arginine rich 1 protein (GAR1), a core small nucleolar ribonucleoprotein (snoRNP) that is an essential component of the mammalian box H/ACA RNP, is recognized as a binding partner of SMN ^12,16^. The H/ACA RNP complex is responsible for catalyzing RNA pseudouridylation in the nucleolus and Cajal bodies, a process crucial for ribosome biogenesis and function ^17–19^. The function of GAR1 has been extensively studied in Archaea, where it facilitates the accurate positioning of substrate RNA and enhances the reaction kinetics within the H/ACA complex ^20–22^. While the archaeal homolog is composed only by a conserved RNA-recognition motif (RRM), eukaryotic GAR1 includes two additional RGG domains at the N- and C-terminus of the core RRM, which may contribute to unidentified GAR1 functions ^23^.

The SMA type I-linked point mutation E134K within the SMN TD has been shown to impair the ability of SMN to interact with the N- and C-terminal RGG domains of GAR1 ^16^. It has been suggested that SMN is involved in the H/ACA RNP assembly, facilitated through its interaction with GAR1 ^12^. In fact, decreased levels of ribosomal RNA (rRNA) pseudouridylation, along with reduced levels of both SMN and H/ACA RNP in Cajal bodies, have been observed in SMA patients, further implicating the role of GAR1 and SMN interaction in SMA pathology ^24–26^.

Recent research has demonstrated that the RGG motifs in yeast homolog GARR-1 contribute to sub-nucleolar condensation ^27^. However, the phase separation of human GAR1 has not been reported to date. Additionally, the potential impact of SMA-associated mutations in the SMN TD on the LLPS behavior of RGG-rich proteins remains unknown.

Here, we demonstrate that the H/ACA snoRNP GAR1 is a scaffold LLPS protein, which undergoes electrostatic-driven coacervation with RNA to form gel-like condensates *in vitro*, and is phase separated in the nucleoli and Cajal bodies of cells. This process is modulated by the interaction between GAR1 and SMN TD, that is disrupted by the SMA-related mutation E134K. In cells, SMN^WT^ co-localized with GAR1 in Cajal bodies, whereas SMN^E134K^ was less abundant in these compartments. Furthermore, we show that altering the stoichiometry of GAR1:RNA affects the distribution of RNA within these condensates, which in turn influences their viscoelastic properties. Using NMR spectroscopy, we demonstrate that SMN TD interacts with GAR1 through its conserved loops, a critical interaction surface that is perturbed by the E134K mutation. Additionally, we show that SMN^E134K^ variant has a moderated impact on GAR1 dynamics *in vitro* and in cells compared to SMN^WT^. *In vitro*, while SMN^WT^ TD alters the morphology of the condensates, promotes local decondensation, and enhances fluidity by competing with RNA for GAR1 binding, these effects are considerably reduced in the E134K variant due to its diminished affinity for GAR1. Our findings suggest a potential LLPS-related mechanism in SMA, centered on the disruption of the interaction between SMN and GAR1.

## Results

### RNA promotes GAR1 phase separation and assembly of gel-like condensates *in vitro*

In order to evaluate the phase separation propensity of human GAR1 protein, we used the sequence-based LLPS predictors FuzDrop and PSPredictor ^28,29^. Both algorithms revealed that GAR1 had a very high probability to phase separate, with a FuzDrop score of P_LLPS_ = 0.998 and a PSPredictor score of PSP = 0.998. FuzDrop suggested that LLPS was mediated by the N-(residues 1-64) and C-terminal (residues 164-217) RGG motifs, with an additional contribution from the RRM region (residues 65-73 and 147-163) **(Fig. 1A**). Additionally, both RGG domains of GAR1 were also predicted to exhibit prion-like characteristics by the PLAAC algorithm, a sequence characteristic of LLPS scaffolds **(Fig. 1B)** ^30,31^.

**Figure 1.**
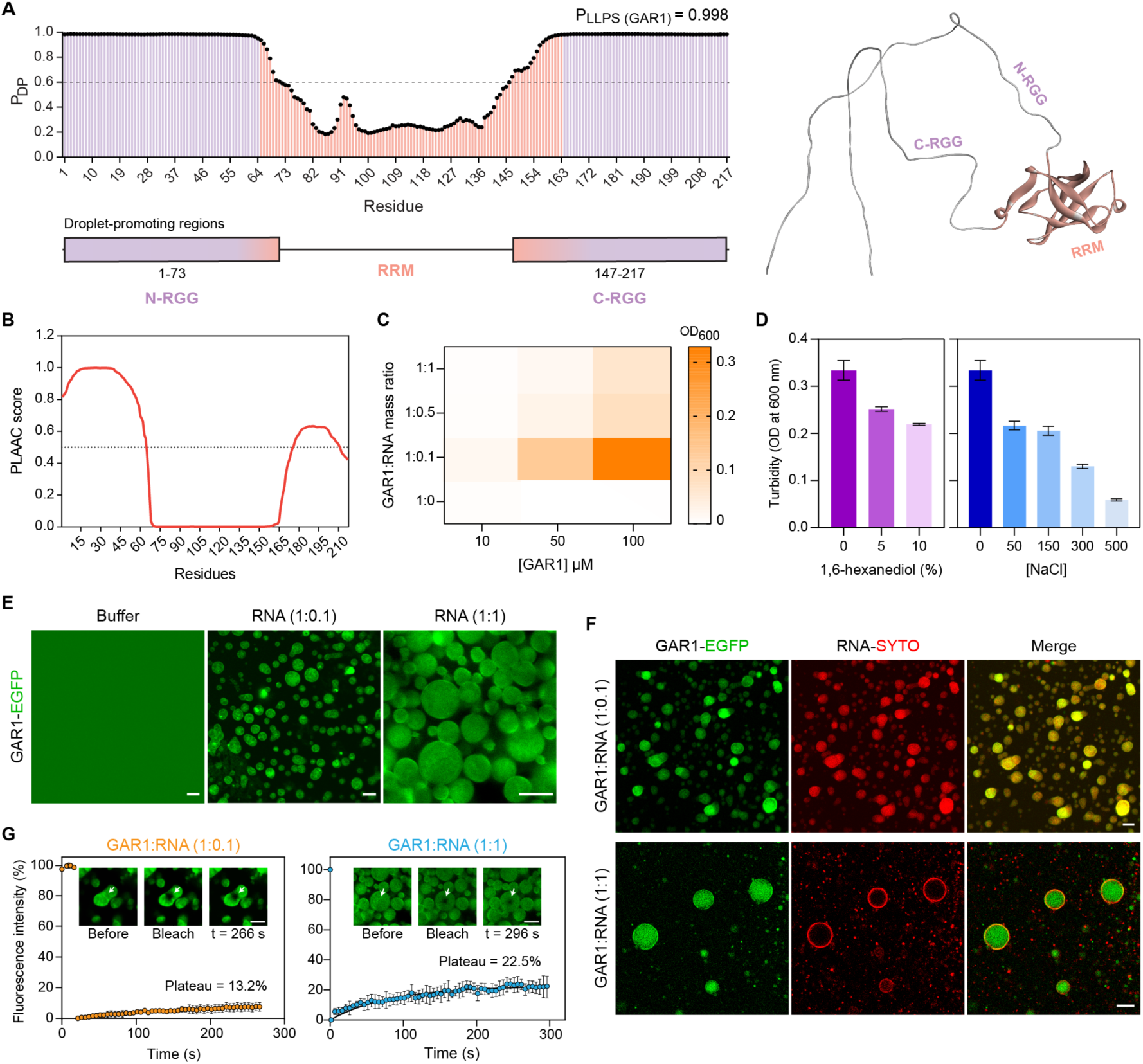
GAR1 protein assembles gel-like condensates through complex coacervation with RNA. **A.** FuzDrop prediction (left panel) estimates that GAR1 phase separation is primarily mediated by the N- and C-terminal RGG domains (in purple). AlphaFold 2 model of human GAR1 structure (UniProt code: Q9NY12) indicating the corresponding protein domains (right panel). **B.** PLAAC algorithm attributes prion-like characteristics to the N- and C-terminal RGG domains. **C.** Phase diagram of GAR1 and RNA at distinct GAR1 concentrations and GAR1:RNA mass ratios. **D.** Turbidity values of 100 µM GAR1 and RNA (1:0.1 mass ratio) in the presence of increasing 1,6-hexanediol and NaCl concentrations. Data are presented as mean ± SD, *n* = 3 measurements per condition. **E.** Confocal microscopy of *in vitro* GAR1 (0.5% GAR1-EGFP) in solution and GAR1-EGFP:RNA condensates. Scale bar, 5 µm. **F.** Confocal microscopy of *in vitro* GAR1 (0.5% GAR1-EGFP) and RNA stained with SYTO dye at different mass ratios. Scale bar, 10 µm. **G.** FRAP analysis of GAR1:RNA condensates at different mass ratios. Inset: confocal image of bleached condensate. Data points are presented as mean ± SD, *n* = 3 independent measurements. P_LLPS_, probability score for spontaneous LLPS; P_DP_, residue-specific probability for droplet formation; error bars, standard deviation (SD); scale bar, 5 µm.

Charge-mediated protein phase separation constitutes a prevalent mechanism governing condensate formation ^32–34^. This type of phase separation involves multivalent electrostatic interactions between oppositely charged polyelectrolytes, including nucleic acids and RGG-rich proteins, and is termed complex coacervation ^32^. Considering the positive net charge of GAR1 at physiological pH (pI = 10.9), mainly clustered in the RGG motifs, and its biological role involving nucleic acid modifications, we sought to investigate the potential complex coacervation of GAR1 and RNA. For this, we conducted turbidity assays and confocal fluorescence microscopy using samples containing GAR1-EGFP:GAR1 in a 1:200 ratio, in the absence and presence of RNA.

In the absence of RNA and under near-neutral pH conditions (6.8), GAR1 samples at distinct concentrations (10, 50 and 100 µM) exhibited no significant turbidity at 600 nm and confocal fluorescence microscopy revealed that GAR1 remained dispersed. Upon adding RNA at a GAR1:RNA mass ratio of 1:0.1, there was a marked increase in turbidity in the 50 µM and 100 µM GAR1 samples, accompanied by the formation of discernible condensates that were detected by microscopy **(Fig. 1C,E)**. The phase diagram, plotted as a function of increasing GAR1:RNA ratio, demonstrated a progressive decrease in turbidity with increasing relative RNA concentration across all tested GAR1 samples, suggesting that the condensates are governed by electrostatic interactions **(Fig. 1C)**. To test this, we monitored the susceptibility of the GAR1:RNA condensates (100 µM in 1:0.1 ratio) to NaCl, which disrupts electrostatic interactions, and 1,6-hexanediol, a compound that dissolves condensates assembled through hydrophobic interactions ^35^ **(Fig. 1D)**. The significant reduction in turbidity with increasing NaCl concentration, coupled with minimal changes following the addition of 1,6-hexanediol, suggests that electrostatic interactions predominantly mediate the coacervation process, while hydrophobic interactions play a minor role **(Fig. 1D)**. Microscopy further reveled that GAR1 condensates exhibit distinct distribution of GAR1-EGFP depending on the GAR1:RNA mass ratio **(Fig. 1E)**. In samples with a GAR1:RNA mass ratio of 1:0.1, the condensates displayed a heterogeneous distribution of GAR1-EGFP within the droplets. Additionally, we observed the presence of species with bright fluorescence and irregular morphology, indicative of potential aggregate formation. In contrast, in GAR1:RNA 1:1 samples, GAR1-EGFP was uniformly distributed, with a reduced incidence of these putative aggregates. To elucidate the co-localization of RNA within the droplets, RNA was stained with SYTO Deep Red Nucleic acid stain. Our results revealed that the distribution of RNA within GAR1 condensates was also dependent on the GAR1:RNA ratio **(Fig. 1F)**. At low RNA levels, RNA was homogeneously distributed, whereas at a 1:1 ratio, RNA preferentially accumulated at the interface of GAR1 condensates. This variation in condensate architecture likely arises from changes in surface tension, viscosity, and polymeric density due to the increased RNA enrichment within the droplets ^31,36,37^.

To investigate the material properties of GAR1 condensates, we performed fluorescence recovery after photobleaching (FRAP) experiments. FRAP analysis revealed that GAR1:RNA condensates displayed gel-like properties, characterized by a limited fluorescence recovery after photobleaching, persisting even after 300 s (a standard time frame typically used to assess liquid-like condensates), by a slow attainment of plateau levels comparable to solid-like assemblies, and by high immobile fractions indicating restricted mobility of fluorescent molecules and low dynamics **(Fig. 1G)** ^38–40^. FRAP analysis allowed us to estimate the apparent diffusion coefficients (*D*_*app*_) for both RNA conditions. Our results revealed that the diffusion coefficient was almost 3-fold higher in GAR1:RNA 1:1 condensates (*D_app_* ≈ (9.32 ± 1.06) x 10^-4^ µm^2^/s) than in 1:0.1 droplets (*D_app_* ≈ (3.33 ± 0.05) x 10^-4^ µm^2^/s). The increased dynamics at higher RNA levels might be attributed to the overall reduction of polymeric density, likely due to the accumulation of RNA at the droplets interface. It is important to note that these *D*_*app*_ values are a thousand-fold lower than those obtained for liquid-like droplets using FRAP techniques, such as FUS condensates (*D_app_* ≈ 0.4 µm^2^/s) and P-granules (*D_app_* ≈ 1 µm^2^/s), suggesting that GAR1:RNA condensates have gel-like characteristics ^38,41^.

These findings demonstrate that GAR1 and RNA undergo complex coacervation *in vitro* leading to the formation of gel-like condensates with viscoelastic properties and droplet architecture influenced by the GAR1:RNA mass ratio.

### GAR1 is phase separated in the cell nucleus

Given that we demonstrated that GAR1 undergoes phase separation *in vitro*, we sought to determine whether this phenomenon also occurred in the cellular environment. To address this, we transfected HeLa cells with EGFP-tagged GAR1 and observed that GAR1 predominantly localizes within the nucleoli, specifically in the dense fibrillar component (DFC) **(Fig. 2A)**. This localization pattern mirrors that of its *Caenorhabditis elegans* homolog GARR-1 ^27^. Additionally, we also observed GAR1 compartmentalization in nuclear bodies and, to a lesser extent, within the granular component (GC) of the nucleolus. This distributions aligns with the known localization of the box H/ACA RNP, which is typically concentrated in the DFC and Cajal bodies ^42^.

**Figure 2.**
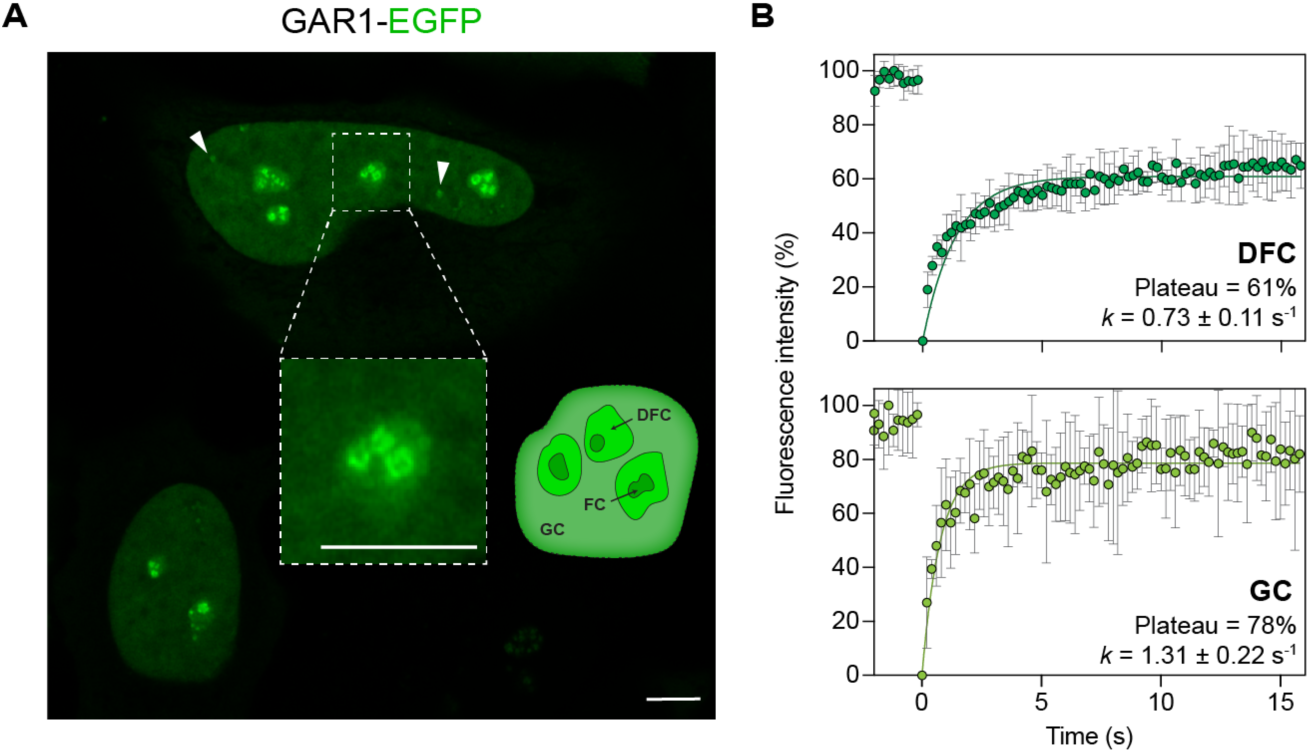
GAR1 localizes to the cell nucleus where it is phase separated in membraneless structures. **A.** Confocal imaging reveals that GAR1 localizes to the nucleolus and nuclear bodies in HeLa cells. White arrowheads indicate the nuclear bodies. **B.** FRAP of different compartments of the nucleolus indicate that GAR1-EGFP has restricted mobility in the DFC compared to in the GC. Data points are presented as mean ± SD, *n* = 5 cells per condition. *k*; recovery rate; DFC, dense fibrillary component; FC, fibrillary component; GC, granular component; scale bar, 5 µm.

To further explore the dynamic properties of GAR1 in the nucleolus, we conducted FRAP analysis of EGFP-GAR1 in both the DFC and GC. The results revealed that while GAR1 displays more liquid-like properties in the GC, accompanied with a fast recovery rate (*k*) and higher diffusion (*D_app_* ≈ 0.70 ± 0.06 µm^2^/s), it is significantly less mobile in the DFC (*D_app_* ≈ 0.33 ± 0.03 µm^2^/s) **(Fig. 2B)**. Although, the apparent diffusion values are still within the liquid-like range, it is important to note that full FRAP recovery is not attained in both compartments, with the recovery being particularly lower in the DFC (plateau of 61%), suggesting that GAR1 has restricted mobility in this sub-compartment. Previous reports have demonstrated that DFC and GC have different viscoelastic properties, with DFC generally exhibiting slower dynamics and less liquid-like characteristics compared to the GC ^36^. These observations not only show that GAR1 is phase separated in the nucleoli but also suggest that GAR1 may contribute to the reduced dynamics of the DFC by promoting gel-like phase separation, as demonstrated by our *in vitro* findings.

### SMA-related mutation E134K in SMN interferes with its localization and modulation of GAR1 dynamics in Cajal bodies

GAR1 has been shown to interact with SMN *in vivo* ^12^, with mutations in the SMN TD impairing this interaction in *in vitro* binding assays ^16^. However, the structural details and functional implications of the GAR1-SMN interaction remain unclear. To explore these implications, we investigated how the prevalent SMA-related mutation E134K in SMN affects GAR1 in cells. For this, we co-transfected HeLa cells with EGFP-tagged GAR1 and either SMN WT or E134K. Our results demonstrated that SMN is localized both in the cytoplasm and in the cell nucleus, and is co-localized with GAR1 in a subset of nuclear bodies. Based on the known subcellular localization of GAR1 and SMN ^12,43–45^, we infer that these GAR1^+^ SMN^+^ nuclear bodies to be Cajal bodies. Compared to SMN^WT^, the E134K mutant was less abundant in the Cajal bodies, while showing identical levels in other cellular compartments **(Fig. 3)**. Given the co-localization of GAR1 and SMN in Cajal bodies, we used Förster resonance energy transfer read out by automated fluorescence lifetime imaging (FLIM-FRET) to assess whether the proteins interact. Our FLIM-FRET data supports the interaction between SMN^WT^ and GAR1 in the Cajal bodies **(Fig. S1A,B)**. Additionally, we did not observe FRET between GAR1 and SMN^E134K^ **(Fig. S1C)**, though difference in signal-to-noise ratio may explain negative results ^46^ **(Fig. S1D)**. However, given the indication that SMN and GAR1 might interact, we used FRAP to evaluate whether this interaction affects GAR1 dynamics in Cajal bodies. FRAP experiments revealed that SMN^WT^ significantly alters the recovery rate of GAR1 after photobleaching in Cajal bodies, while SMN^E134K^ failed to do so **(Fig. 4)**. Values of recovery rates in the presence of SMN^E134K^ are similar to those of GAR1 alone, suggesting that the GAR1-SMN^WT^ interaction is critical to modulate GAR1 dynamics in Cajal bodies.

**Figure 3.**
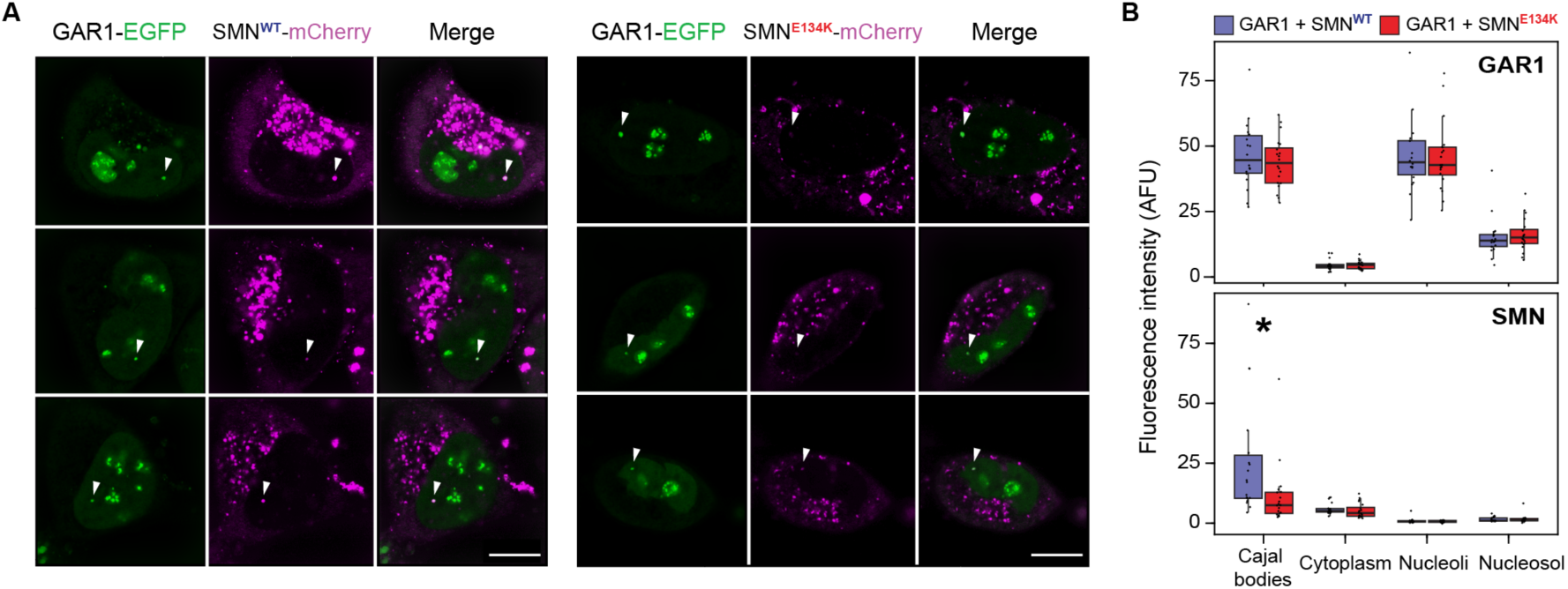
The SMA-related E134K mutation in SMN reduces its localization to Cajal bodies. **A.** Representative images of HeLa cells co-transfected with plasmid DNA encoding GAR1-EGFP and mCherry-SMN (WT or E134K mutant), at 4 days post-transfection. Arrowheads point to Cajal Bodies, where GAR1 and SMN colocalize. **B**. Mean background-corrected fluorescence intensity of GAR1-EGFP or mCherry-SMN in different cellular compartments. Data points are presented as mean ± SD, *n* = 10-20 cells per condition. * P < 0.05, mCherry-SMN intensity in Cajal bodies of cells co-transfected with GAR1 + SMN^WT^ vs. GAR1 + SMN^E134K^; ANOVA with Tukey’s post-hoc on log-transformed data; scale bar, 10 µm.

**Figure 4.**
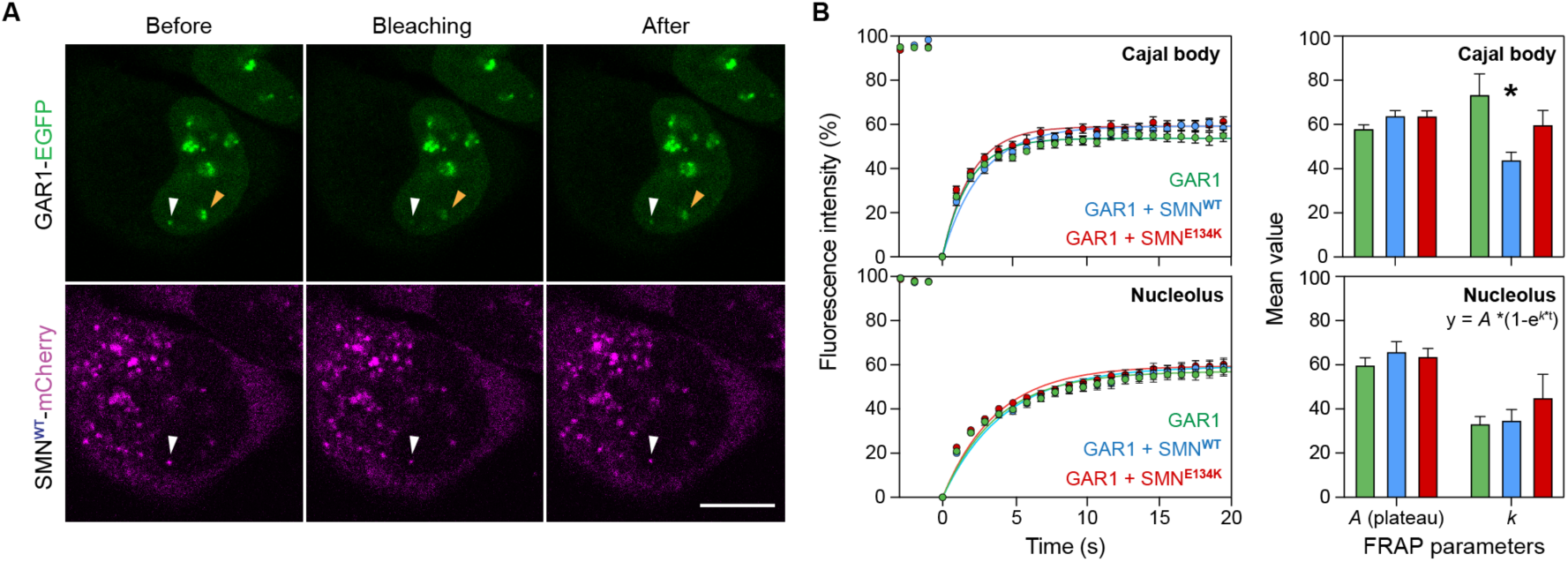
Wild-type but not mutant SMN alters GAR1 dynamics in Cajal bodies. Bleaching was performed with a 488 nm laser line on Cajal bodies and nucleoli of HeLa cells co-transfected with GAR1 + SMN. **A.** Representative images of fluorescence recovery after photobleaching (FRAP); white and yellow arrowheads point to Cajal body and nucleolus, respectively. **B.** FRAP quantifications of cells transfected with either GAR1 alone, GAR1 + SMN^WT^, or GAR1 + SMN^E134K^, on bleached Cajal bodies (upper panel) and nucleoli (lower panel). FRAP curves, normalised fluorescence intensity vs. time (left panel). Parameter values of single exponential decay models fit to FRAP curves (right panel). Data points are presented as mean ± SD, *n* = 53-54 Cajal bodies or nucleoli per condition. * P < 0.05, k value in Cajal bodies of cells transfected with GAR1 + SMN^WT^ vs. GAR1 + SMN^E134K^, ANOVA with Tukey’s post-hoc on log-transformed data. *k*; recovery rate.

### The SMN Tudor domain interacts with GAR1 through its conserved loops

Since our FLIM-FRET data in cells suggested an interaction between GAR1 and SMN within the Cajal bodies, we aimed to investigate this interaction in detail. For this, we focused on the TD of SMN, since it is known to mediate this binding ^16^.

Through microscale thermophoresis (MST), we determined that SMN^WT^ TD (residues 83-147) has a micromolar affinity for GAR1 with a dissociation constant (KD) of 8 ± 1 µM **(Fig. S2A)**. Then, using solution-state Nuclear Magnetic Resonance (NMR) spectroscopy, we successfully transferred and confirmed the backbone and side-chain chemical shifts of SMN^WT^ TD using the assignment data deposited in the Biological Magnetic Resonance Data Bank (BMRB:4899) by Selenko *et al.* ^14^ **(Fig. S2B)**. To map the intermolecular interaction between SMN^WT^ TD and GAR1, we performed NMR titration experiments. For this, ^15^N-labeled SMN^WT^ TD with a fixed concentration of 100 µM, was titrated with increasing concentrations of unlabeled GAR1 and ^1^H-^15^N heteronuclear single quantum coherence (HSQC) spectra were acquired. The titration points corresponded to SMN^WT^ TD:GAR1 concentration ratios of 1:0.25, 1:0.50, 1:0.75, and 1:1. As a preliminary control to these experiments, we verified through turbidity assays and confocal fluorescence microscopy that SMN^WT^ TD does not promote significant droplet formation of GAR1 under our experimental conditions, which could interfere with the interaction characterization **(Fig. 5A)**. As an additional control, we acquired an ^1^H-^15^N HSQC spectrum of the 1:1 titration point with 10 µM of GAR1/SMN TD, where phase separation is negligible, which showed that the same residues were affected when compared with the identical titration point with 100 µM. This confirmed that LLPS does not affect the results **(Fig. 5A and S2C)**. The titration experiments revealed that the interaction between the SMN^WT^ TD and GAR1 primarily occurs via the conserved loops 1 and 3 of SMN TD **(Fig. 5B and S2D)**, on a fast chemical exchange timescale **(Fig. 5C)**. Loop 1 contains the residues Glu104 and Asp105, which form a negatively charged surface that likely interacts with the positively charged arginine residues of GAR1 **(Fig. 5D)**. Within this loop, and including some residues of the β1 strand, we observed significant chemical shifts in residues from Ala100 to Tyr109, except Trp102, which is buried in the hydrophobic core of SMN^WT^ TD ^14^. These findings align with previous reports demonstrating that missense mutations within loop 1 reduced GAR1 binding to SMN TD by 70-95% ^16^. Additionally, significant chemical shift perturbations were encountered in residues belonging to loop 3, spanning from Thr128 to Arg133. This region features a hydrophilic patch, characterized by the presence of the conserved residues Tyr130 and Gly131 **(Fig. 5D)** ^14^. Together, these results support the findings reported by Selenko *et al*., demonstrating that the Sm D_1_ protein binds to the loops 1 and 3 of SMN^WT^ TD ^14^. Given that the Sm D_1_ protein contains a C-terminal RG domain similar to that of GAR1, our data supports that GAR1 interacts with SMN^WT^ TD loops through the RGG domains, as previously proposed by S. Whitehead *et al* ^16^.

**Figure 5.**
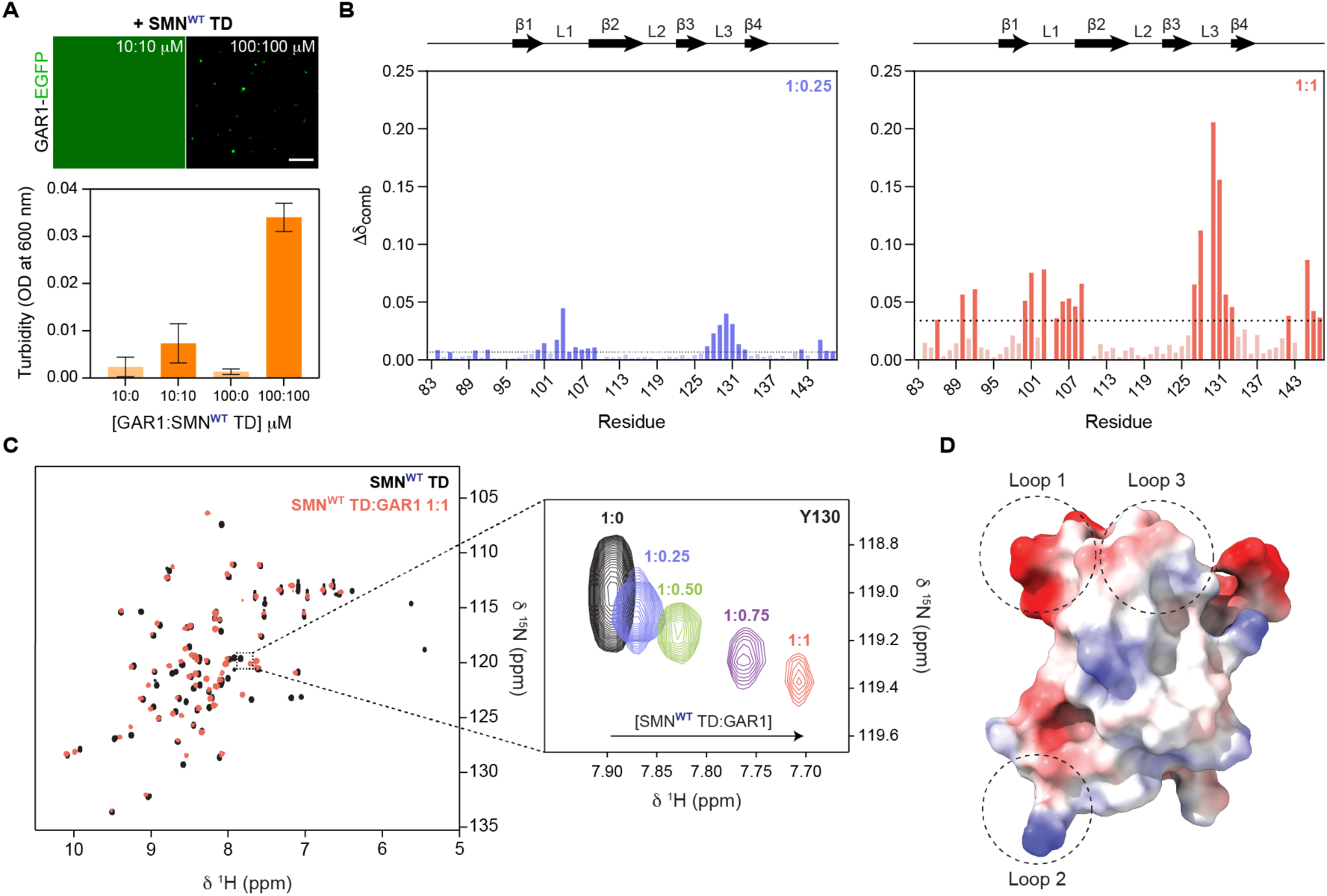
GAR1 interacts with conserved loops in the Tudor domain of SMN. **A.** Confocal microscopy and turbidity measurements of GAR1 and SMN TD at distinct concentrations in equimolar ratios. Data points are presented as mean ± SD, *n* = 3 independent measurements. Scale bar, 5 µm. **B.** Combined chemical shift perturbation of ^15^N-labeled SMN^WT^ TD resonances with the addition unlabeled GAR1, at 1:0.25 (left panel, blue) and 1:1 (right panel, red) molar ratios. Bars in bold color indicate the estimated interacting residues (above the cut-off). Above the plots, a visual representation of the secondary structure of SMN TD is depicted. **C.** Overlay of ^1^H-^15^N HSQC spectra of ^15^N-labeled SMN^WT^ TD alone (black) and in the presence of unlabeled GAR1 in a 1:1 molar ratio (red). Example of chemical shift perturbation of Y130 residue during titration with unlabeled GAR1. **D.** Electrostatic surface representation of the AlphaFold 2 model of SMN^WT^ TD construct used in this study (UniProt code: Q16637, fragment 83-147), indicating the loops location. Blue and red color depict the positive and negative electrostatic surface potential with coulombic values from - 10.45 to +12.15, respectively. β – beta-sheet, L – loop.

### SMA-related mutation E134K in the SMN Tudor domain interferes with the interaction with GAR1 through its binding loops

To investigate the impact of the SMA-related TD pathological mutation E134K on its interaction with GAR1, we conducted similar experiments to those previously performed for SMN^WT^ TD. Consistent with previous reports, we detected that the SMN TD point mutation E134K resulted in a substantial decrease in affinity for GAR1, which precluded the determination of binding affinity values via MST **(Fig. S3A)** ^16^. Contrary to SMN^WT^ TD, no assignment data for the mutated E134K variant is available in the BMRB. As a result, we initially attempted to assign SMN^E134K^ TD based on the assignment for SMN^WT^ TD. However, given the significant differences observed in the ^1^H-^15^N HSQC spectra between the two variants, additional 3D NMR experiments were required to validate the final assignment. We successfully assigned all SMN^E134K^ TD backbone ^1^H-^15^N resonances except for Lys97, Ser103 and Glu104 **(Fig. S3B)**. In addition, {^1^H}-^15^N heteronuclear nuclear Overhauser effect (hetNOE) experiments, which measure hetNOE ratios that are correlated with the mobility of each amide in the protein backbone ^47^, indicated that the E134K mutation does not disrupt the overall ns/ps dynamics of the TD **(Fig. S3C)**. This suggests that the folded structure of the SMN TD remains intact, which is in agreement with previous reports by Selenko *et al* ^14,48^.

To map the interaction between SMN^E134K^ TD and GAR1, we performed NMR titration experiments similar to those conducted with SMN^WT^ TD. Titration points corresponded to SMN^E134K^ TD:GAR1 ratios of 1:0, 1:0.25, 1:0.50 and 1:1. As a control, we also confirmed that the E134K variant does not induce significant LLPS of GAR1 **(Fig. 6A)**. Given that the turbidity values and presence of droplets were even lower than those observed with SMN^WT^ TD, and considering our prior demonstration that the minimal droplet formation with SMN^WT^ TD did not affect the titration results **(Fig. S2C)**, we assumed that the presence of E134K also did not influence the results. The NMR experiments captured interactions between the two proteins and demonstrated that they are mediated through the same regions of the SMN^WT^ TD, specifically loop 1 and loop 3 **(Fig. 6B and S3D,E)**. As expected, however, the interactions were less pronounced compared to those between SMN^WT^ TD and GAR1, particularly those involving loop 3. Specifically, the chemical shift perturbations in residues within loop 3, such as Thr128, Tyr130, and Gly131, were on average 8.5-fold lower relative to SMN^WT^ TD. In contrast, the chemical shift perturbation in residues within loop 1 was approximately 1.5-fold lower when compared to SMN^WT^ TD. These findings show that while the E134K mutation weakens the overall interaction with GAR1, the most pronounced impact is observed in interactions involving loop 3. These results provide important residue-level information consistent with previous pull-down assays findings by Whitehead *et al* ^16^. Although Lys134 may not directly participate in interactions with GAR1 (as its resonances do not undergo significant chemical shift perturbation), the inversion of electrostatic potential in loop 3, due to the mutation, might impact interactions of nearby residues **(Fig. 6C)**. As previously stated, our results coupled with published data suggest that SMN TD interacts with the prion-like RGG domains of GAR1 ^14,16^. Since the arginine residues of GAR1 are positively charged at the experimental pH (6.8), due to the high pK_a_ (12.5) of the guanidinium group present on their side-chains, the positive charge of the lysine of SMN^E134K^ TD could disrupt nearby interactions in loop 3 through electrostatic repulsion, particularly affecting the cation-ρε interactions with TD’s aromatic residues, such as Tyr130.

**Figure 6.**
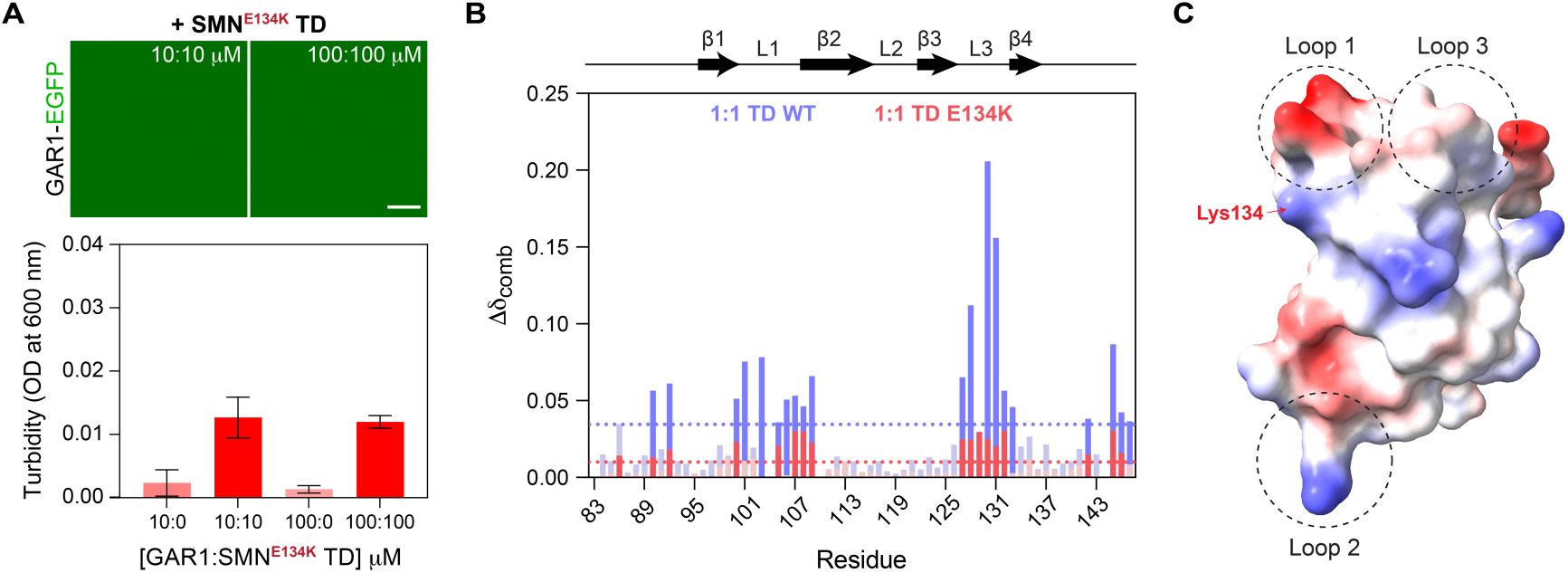
The E134K mutation in the Tudor domain of SMN compromises its interaction with GAR1. **A**. Confocal microscopy and turbidity measurements of GAR1 and SMN^E134K^ TD at distinct concentrations in equimolar ratios. Data points are presented as mean ± SD, *n* = 3 independent measurements. Scale bar, 5 µm. **B.** Overlay of combined chemical shift perturbation of ^15^N-labeled SMN TD WT (purple) and E134K (red) resonances in the 1:1 equimolar ratio of SMN TD:GAR1. Bars in bold color indicate the estimated interacting residues (above the indicated cut-off). Above the plots, a visual representation of the secondary structure of SMN TD is depicted. **C.** Electrostatic surface representation of the AlphaFold 2 model of SMN^E134K^ TD construct used in this study (UniProt code: Q16637, fragment 83-147 K134), indicating the loops and mutated lysine location. Blue and red color depict the positive and negative electrostatic surface potential with coulombic values from -11.37 to +10.22, respectively. β – beta-sheet, L – loop.

### SMN TD WT drives reentrant phase separation in GAR1:RNA condensates through competitive RNA interactions

Next, we aimed to explore and characterize the molecular and biophysical consequences of the presence of SMN^WT^ or SMN^E134K^ on GAR1 condensates. For this, we established a simplified model of reconstituted condensates, as *in vitro* mimics of Cajal bodies, using recombinant GAR1, SMN TD, and RNA. First, we confirmed that SMN TD did not undergo LLPS under our experimental conditions **(Fig. S4A,B)**. Next, we examined the localization of SMN TD WT and E134K variant within GAR1:RNA condensates, at both RNA ratios studied (1:0.1 and 1:1), by labeling the TDs with Alexa-Fluor 647. Confocal fluorescence images revealed that both SMN TD WT and E134K partitioned to the droplets exclusively when RNA was present at a GAR1:RNA ratio of 1:0.1 **(Fig. 7A)**. In this condition, it was possible to observe that SMN^WT^ TD promoted reentrant phase separation, inducing the formation of vacuoles within the GAR1:RNA condensates, that effectively excluded GAR1-EGFP and SMN^WT^ TD **(Fig. 7A,B)**. Previous studies in different systems have also shown the formation of such aqueous-phase vesicle-like structures in the presence of RNA ^49,50^. The addition of RNA at a 1:1 GAR1:RNA mass ratio resulted in the displacement of SMN^WT^ TD from the condensates **(Fig. 7A)**. These results indicate that there is a competition between SMN^WT^ TD and RNA for GAR1 binding, driving local decondensation and displacing SMN TD as RNA levels increase. To test this competition, we performed NMR experiments of ^15^N SMN^WT^ TD and GAR1 in a 1:1 molar ratio at 10 µM (to avoid RNA-promoted LLPS), while increasing the relative RNA mass. We observed that the chemical shifts of SMN^WT^ TD returned to their original values as RNA was progressively added, giving evidence of this competitive interplay **(Fig. 7C)**. Due to SMN^E134K^ TD’s weaker binding, it was not possible to perform the NMR competition experiments. To monitor the binding to SMN^E134K^ TD, we would need higher concentrations of GAR1, which would phase separate upon addition of RNA.

**Figure 7.**
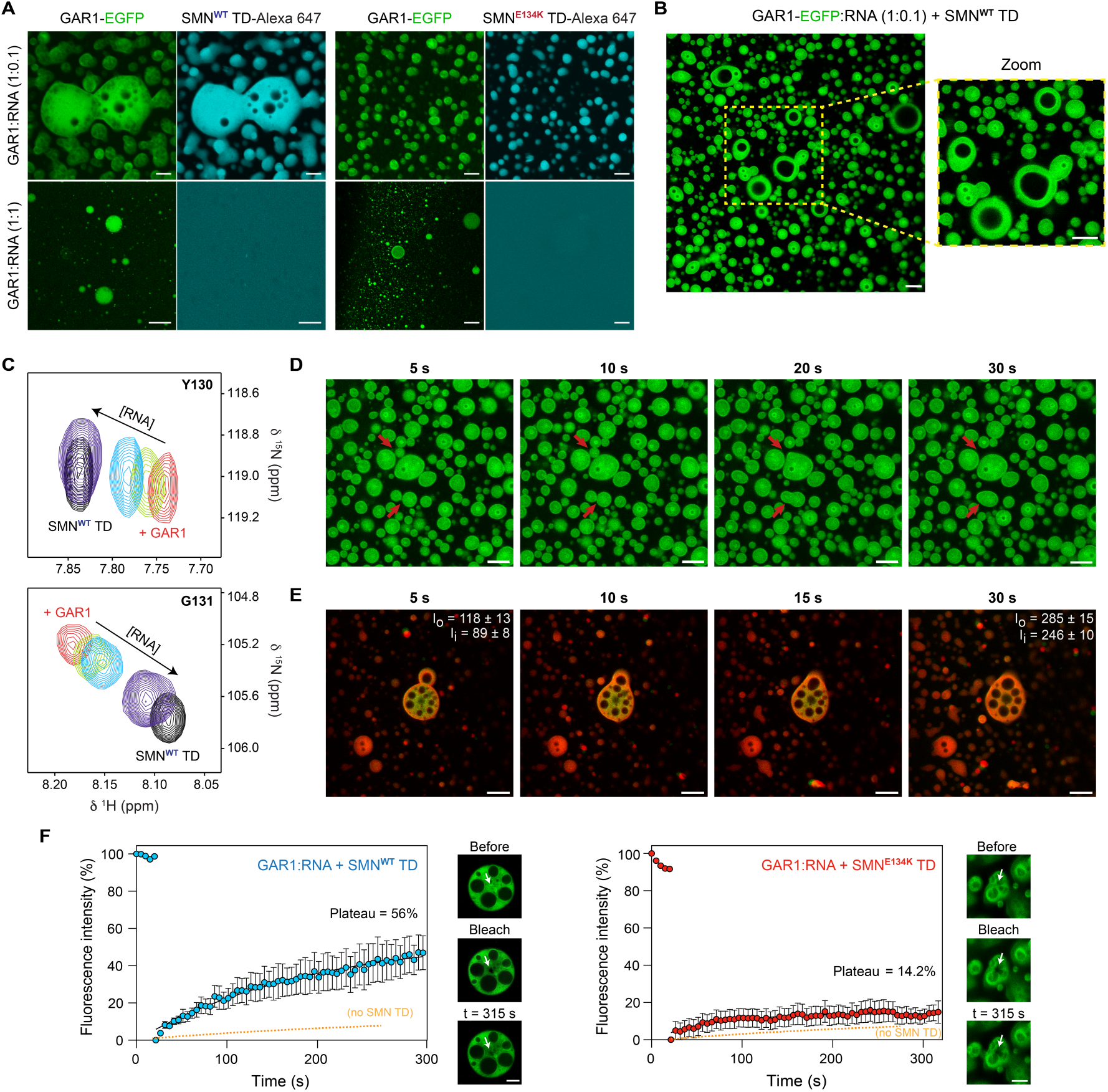
Competition between RNA and SMN Tudor modulates the morphology and dynamics of GAR1:RNA condensates. **A.** Confocal *in vitro* imaging of the co-localization of 100 µM SMN TD WT and E134K (1% SMN TD-Alexa Fluor 647) in 100 µM GAR1:RNA (0.5% GAR1-EGFP) condensates at low GAR1:RNA ratios, and partition inhibition at 1:1 mass ratios. **B.** Vacuole formation in GAR1:RNA droplets induced by SMN^WT^ TD. **C.** NMR competition experiments showing that RNA displaces GAR1 from the interaction with SMN TD. Examples of 10 µM ^15^N-labeled SMN^WT^ TD ^1^H-^15^N HSQC chemical shifts, belonging to Y130 and G131 residues, returning to initial values as RNA is added in increasing amounts (black – SMN^WT^ TD, red – addition of unlabeled GAR1 in a 1:1 SMN TD:GAR1 ratio, green – addition of RNA in a 1:0.1 GAR1:RNA ratio, light blue – 1:0.5, dark blue – 1:1). **D.** Time-lapse series showing *in vitro* GAR1 and RNA condensates with SMN^WT^ TD (1:0.1:1 ratio) fusing over time. **E.** Time-lapse series of *in vitro* GAR1 and RNA (SYTO stain) and SMN^WT^ TD (1:0.1:1 ratio) demonstrating the localization of RNA at the droplet boundaries and the merging of vacuoles. SYTO intensity at the outer edge (I_o_) and inside (I_i_) the droplets equalize overtime, indicating dispersion of RNA molecules as the fusion events occurs. **F.** FRAP analysis of GAR1:RNA 1:0.1 condensates with SMN TD WT (left panel) and E134K (right panel). FRAP fitting of the GAR1:RNA 1:0.1 (no SMN TD) condition is plotted for comparison (dashed orange line). Data points are presented as mean ± SD, *n* = 3 independent measurements. Insets: confocal image of bleached condensates. Scale bars, 5 µm.

Additionally, we observed that these multiphasic condensates remained fluid and fusion events occurred **(Fig. 7D)**. By staining the RNA with SYTO dye, we could observe that the RNA was also excluded from the internal vacuoles observed in the droplets, further confirming their hollow/aqueous nature **(Fig. 7E)**. During fusion events, these hollow droplets became trapped within larger condensates and fused internally with additional cavities, resulting in the formation of larger vacuoles **(Fig. 7E)**. RNA molecules also dispersed around the condensate during fusion events, suggesting their active involvement in this fusion process and subsequent rearrangement within the resulting structure. Although equally excluded from the droplets upon adding RNA, SMN^E134K^ TD variant failed to induce discernible vacuoles on GAR1:RNA condensates **(Fig. 7A)**. Since SMN^E134K^ TD affinity for GAR1 is weaker in comparison to WT TD, it might fail to outcompete RNA for GAR1 binding and promote reentrant phase separation.

### The E134K mutation impairs the ability of the SMN Tudor domain to fluidize the GAR1:RNA condensates

After assessing the distinct effects of SMN TD WT and the E134K variant on GAR1:RNA condensate morphologies, we sought to examine their influence on condensate dynamics. FRAP results revealed that SMN^WT^ TD significantly increased the dynamics of GAR1:RNA condensates, while SMN^E134K^ TD exhibited a reduced impact **(Fig. 7F)**. Notably, the WT variant decreased the immobile fraction from 87 ± 1.45% (in the absence of SMN TD) to 44 ± 1.86% within the same time frame. Additionally, the apparent diffusion coefficient for GAR1:RNA 1:0.1 condensates in the presence of SMN^WT^ TD was double (*D_app_* ≈ (7.61 ± 0.45) x 10^-4^ µm^2^/s) that of GAR1:RNA 1:0.1 (*D_app_* ≈ (3.33 ± 0.05) x 10^-4^ µm^2^/s). Despite the two-fold difference in apparent diffusion coefficients, both values remained within the standard values for gel-like condensates ^51,52^.

In contrast, the introduction of SMN^E134K^ TD did not result in significant changes in the immobile fraction, remaining at to 86 ± 0.28% compared to 87 ± 1.45% (in the absence of SMN TD), and led to a modest 1.3-fold increase in the diffusion coefficient (*D*_*app*_≈ (4.50 ± 0.3) x 10^-4^ µm^2^/s). This discrepancy in the modulation of GAR1:RNA condensate dynamics may stem from the distinct affinities of the SMN TD variants for GAR1. While SMN^WT^ TD efficiently displaces RNA molecules from GAR1, thereby increasing condensate fluidity, SMN^E134K^ TD’s reduced affinity for GAR1 limits its ability to compete significantly with RNA, resulting in a smaller effect on GAR1:RNA condensate dynamics.

## Discussion

In human neurodegenerative conditions, accumulating evidence has linked pathological mutations of scaffold RNA-binding proteins to the dysfunction of condensates ^39,53–55^. However, the impact of mutations in co-scaffolds and client proteins in these processes remains poorly understood. In this study, we investigated the *in vitro* and cellular phase separation of the eukaryotic H/ACA snoRNP GAR1, revealing that GAR1 undergoes electrostatically-driven coacervation with RNA to assemble gel-like condensates. The modulation of GAR1 phase separation involves a delicate balance of RNA distribution, since increased RNA levels resulted in their accumulation at the condensate interface, impacting condensate architecture and dynamics. This regulation may result from the controlled distribution of RNA within the droplets, a phenomenon that has been previously demonstrated for tau:PrP heterotypic assemblies ^32^. Based on these observations, we hypothesize that GAR1 phase separation may contribute to the accumulation of RNA after its catalytic modification within the eukaryotic H/ACA snoRNP complex, however, further studies are required to test this hypothesis.

Our findings further indicate that, in cells, GAR1 is phase-separated within Cajal bodies and nucleoli, where it exhibits constrained dynamics. Supported by our *in vitro* results, these observations suggest that GAR1 may contribute to the reduced liquid-like environment of these sub-compartments, potentially playing a role in their spatial organization and functional specialization, as observed for its yeast homolog ^27^.

Nevertheless, an important question remained: how could modified RNAs be released from the GAR1 condensates to fulfill their biological functions? Our data indicates that the SMA-associated protein SMN co-localizes and interacts with GAR1 within Cajal bodies, altering its dynamics and potentially regulating RNA release from GAR1 condensates. We show that the TD of SMN acts as a client for GAR1 condensates by binding to GAR1 and competing with RNA, which could drive condensate disassembly and facilitate RNA release. This mechanism was supported by NMR spectroscopy and confocal microscopy data, which revealed that SMN TD induces local decondensation in GAR1:RNA condensates, and that increasing RNA levels weakens GAR1-SMN TD interaction, leading to enhanced condensate fluidity.

Strikingly, the SMA-related E134K mutation in the SMN TD significantly impairs its interaction with GAR1. Although SMN^E134K^ still partitions into GAR1:RNA condensates, its unable to promote local decondensation, and its effects on GAR1 dynamics *in vitro* and within the Cajal bodies are notably reduced. This finding suggests that the E134K mutation disrupts SMN’s ability to regulate RNA release from GAR1 condensates, likely due to its reduced affinity for GAR1.

We hypothesize that the pathological mutation in SMN TD impairs the modulation of GAR1 phase separation, particularly by disrupting RNA release, which may hinder proper RNA processing and functions, contributing to the cellular defects observed in SMA **(Fig. 8)**. This hypothesis aligns with evidence linking defects in rRNA processing, ribosomal depletion, and translational dysfunction to SMA pathology ^56,57^. While further investigations are required to fully elucidate the implications of these findings, our work highlights the importance of SMN-dependent regulation of GAR1 condensates and suggests that targeting this interaction could provide a novel therapeutic strategy for SMA-related dysfunctions.

**Figure 8.**
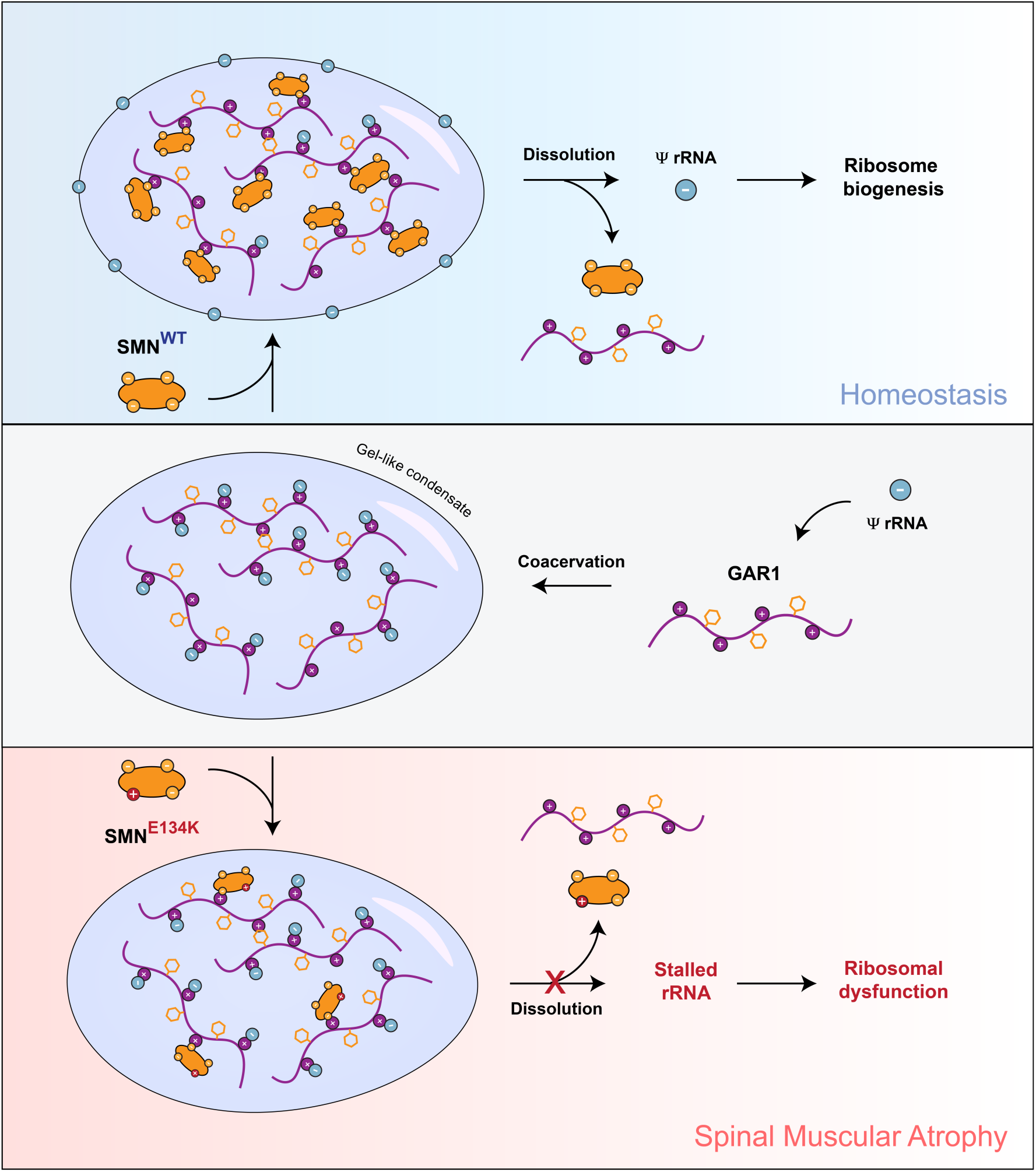
Schematic hypothesis of the SMN-dependent modulation of GAR1 condensates in homeostasis and SMA pathology. Under physiological conditions, GAR1 undergoes complex coacervation, facilitating the release and accumulation of pseudouridine-modified ribosomal RNA0 (ψ rRNA) from the H/ACA complex. SMN binds to GAR1 and displaces ψ rRNA from the condensates through competitive interactions, allowing the RNA to participate in ribosome biogenesis. In SMA, the E134K mutation in SMN impairs its interaction with GAR1, preventing the release of ψ rRNA from the condensates. Consequently, modified ψ rRNA may become stalled, contributing to ribosomal dysfunction observed in SMA.

## Methods

### Prediction of LLPS propensity and prion-like regions

Phase separation predictions were conducted using the FuzDrop web server (https://fuzdrop.bio.unipd.it/predictor), which performs a sequence-based identification of both droplet-promoting and aggregation hotspots based on the query sequence. The FuzDrop output provides a probability score of spontaneous LLPS (P_LLPS_ ≥ 0.6 for scaffolds) and an interactive graph displaying the droplet-promoting probabilities (pDP) per residue ^28,58^. The LLPS propensity was also analyzed through the PSPredictor (http://www.pkumdl.cn:8000/PSPredictor/), a tool that predicts potential phase separation of proteins (PSPs) based on the amino acid sequence through machine learning. The output assigns a PSP score between 0 and 1, and the query protein is classified as a phase-separating protein if its PSP score is ≥ 0.5 ^29^. The propensity of phase separation was analyzed for human GAR1 (UniProt code: Q9NY12) and for SMN TD WT and E134K variant (UniProt code: Q16637; residues 83-147). The prion-like propensity of GAR1 domains was predicted through the PLAAC (Prion-like amino acid composition) algorithm, using default parameters and a relative weighting of background frequencies set at 50%. Regions are considered to be prion-like if the PLAAC score is ≥ 0.5 ^30^.

### Recombinant protein production

Four recombinant proteins were produced in this work, including full-length human GAR1, GAR1 fused with C-terminal enhanced GFP (GAR1-EGFP), wild-type SMN TD, and E134K mutant SMN TD. To facilitate the expression and purification, all recombinant proteins included a N-terminal maltose-binding protein (MBP) tag comprising a 6x poly-histidine sequence at the N-terminus and a tobacco etch virus (TEV) cleavage site at the C-terminus. The amino acid sequence of each protein construct is available in **Table S1**. pHTMP vectors containing the cDNA corresponding to each construct were acquired from NZYTech. Each construct was expressed in *Escherichia coli* BL21(DE3) cells in LB for unlabeled and M9 minimal media for isotopically-labeled proteins, induced at mid-exponential growth with 0.5 mM IPTG for 4h at 37°C (for SMN TD constructs) or overnight at 25°C (for GAR1 constructs).

Harvested cell pellets were resuspended in lysis buffer (for GAR1 proteins: 50 mM Tris-HCl, 1 M NaCl, 10% glycerol, 1 M urea, 50 mM glycine, 10 mM imidazole, 2 mM β-mercaptoethanol, 0.1 mM PMSF, 0.03% NaN_3_, pH 7.4; for SMN TD proteins: 50 mM Tris-HCl, 0.5 M NaCl, 10 mM imidazole, 2 mM β-mercaptoethanol, 0.1 mM PMSF, 0.03% NaN_3_, at pH 7.4) supplemented with an EDTA-free complete mini protease inhibitor tablet (Roche), and lysed through sonication. The soluble extracts were isolated by centrifugation and the clear lysates were loaded onto a HisTrap FF Crude column (Cytiva). The proteins were eluted with 250 mM imidazole, followed by the cleavage of the MBP-tag. TEV protease was added to the protein fractions, and digestion was carried out overnight in parallel with dialysis to remove the imidazole. Cleaved products were loaded onto HisTrap HP column (Cytiva) and eluted in the flow-through. The purified samples, as confirmed by SDS-PAGE, were buffer exchanged to the experimental buffer (20 mM HEPES, 1 mM DTT, 0.03% NaN_3_, pH 6.7), concentrated using VivaSpin Turbo 15 (3 or 10 kDa MWCO) ultrafiltration units (Sartorius), flash-frozen and stored at -80°C until further use.

SMN TD constructs were fluorescently labeled with Alexa Fluor 647 Cys-maleimide (Invitrogen), according to the manufacturer’s procedure. Briefly, proteins were extensively washed using ultrafiltration units to remove DTT. Subsequently, tris(2-carboxyethyl)phosphine (TCEP) was added to the samples to a final concentration of 1 mM. Alexa Fluor 647 was added to the protein samples in a 5x molar excess and incubated overnight at 4°C. Unreacted fluorophore was removed using VivaSpin 3 kDa ultrafiltration units (Sartorius). The labeling efficiency was estimated to be between 1 to 1.5 moles of dye per mole of protein.

### HeLa cell maintenance

HeLa cells have been previously used to study SMN ^59,60^, given their advantageous features of high transfection efficiency, flat morphology (useful for imaging), and prominence Cajal bodies ^61^. Here, HeLa cells (ATCC) were cultured in high-glucose Dulbecco’s Modified Eagle’s Medium with GlutaMAX and pyruvate (DMEM), supplemented with 10% fetal bovine serum (FBS) and antibiotic-antimycotic solution (100 U/mL penicillin, 100 μg/mL streptomycin, and 0.25 μg/mL amphotericin B) (Gibco, ThermoFisher Scientific, Paisley, United Kingdom). Cells were maintained at 37 °C, in humidified air with 5% CO_2_.

### Transient transfection of GAR1-EGFP and mCherry-SMN

To express the proteins of interest, HeLa cells were transfected with plasmids encoding either GAR1-EGFP, mCherry-SMN^WT^, or mCherry-SMN^E134K^, under control of the CMV promoter and enhancer (Genscript, Rijswijk, Netherlands). The amino acid sequence of each protein construct is available in **Table S2**. HeLa cells were seeded at 4.5×10^4^ cells/cm^2^ in 8-well Nunc Lab-Tek chambered coverglass (ThermoFisher). 24 hours after seeding, cells were transfected with Lipofectamine 3000 (ThermoFisher), according to manufacturer’s instructions. Briefly, for each well (0.8 cm^2^), 250 ng plasmid DNA were mixed with 0.375 μL Lipofectamine 3000 reagent and 0.5 μL P3000 reagent in 25 μL Opti-MEM reduced serum medium (ThermoFisher). The resulting mixture was incubated for 5 minutes at room temperature and added to the culture medium. This medium was replaced with fresh culture medium 16 hours after transfection. Imaging assays were performed 1–5 days after transfection.

### Live cell imaging

HeLa cells were imaged in phenol red-free DMEM supplemented with HEPES buffer (ThermoFisher), at 37 °C, using a ZEISS LSM 880 confocal equipped with a 63× 1.4 NA DIC oil immersion objective (Zeiss, Oberkochen, Germany). A 488 nm Argon laser was used to excite EGFP, and a 561 nm diode-pumped solid-state laser was used to excite mCherry. For high-resolution imaging, the Airyscan detector was used in super-resolution mode. For FRAP experiments, a gallium arsenide phosphide (GaAsP) detector was used. The Zeiss Zen Black software was used to control the microscope and to process the raw Airyscan images.

### FLIM-FRET measurements

Fluorescence lifetime imaging was performed with single-photon excitation on a multimodal time-resolved fluorescence microscope. This combined an 80 MHz, near-infrared, femtosecond excitation source (Insight X3, Spectra Physics, Crewe, UK), second harmonic generation unit (Harmonixx SHG, APE, Berlin, Germany), DCS-120 laser scanning unit (Becker & Hickl, Berlin, Germany), inverted microscope (Axio Observer 7, Zeiss, Cambridge, UK) with high (1.4) numerical aperture objective (Plan-Apochromat 63x/1.4 Oil M27, Zeiss, Cambridge, UK), two ultrafast hybrid detectors (HPM-100-07, Becker & Hickl, Berlin, Germany) and time-correlated single photon counting (TCSPC) electronics (SPC-180NX, Becker & Hickl, Berlin, Germany). Images were acquired for 3 minutes per field of view with 423 nm excitation, to minimize the ratio of acceptor to donor excitation, and 500–540 nm emission filtering. Photon counts were histogrammed at 14.6ps time intervals and curve fitting analysis was performed in SPCImage (Becker & Hickl, Berlin, Germany).

### Microscale thermophoresis (MST)

MST experiments were performed at 25°C using a Monolith NT.115 instrument (NanoTemper Technologies) with an MST power of 40% and excitation of 20%. Sixteen samples were prepared and loaded onto capillaries, containing Alexa Fluor 647-labeled SMN TD WT or E134K variant (target) and unlabeled GAR1 (ligand) in 20 mM HEPES, 1 mM DTT, 0.03% NaN_3_, pH 6.8 (experiment buffer) with 0.05% Tween. The concentration of the target was kept constant at 10 nM for SMN^WT^ TD and 20 nM for SMN^E134K^, while the ligand concentration ranged from 8 nM to 260 μM between capillaries. MO. Affinity Analysis software (NanoTemper Technologies) was used to analyze and fit binding equations to the data from three biological replicates, for each SMN TD variant, to determine the K*_D_* values ^62^.

### Turbidity measurements

Turbidity of protein samples was estimated from the optical density at 600 nm, recorded on a SpectraMax microplate reader (Molecular Devices) using SpectraDrop low volume 24-well plates (Molecular Devices). All measurements were carried out at 25°C. Turbidity was monitored 30 minutes after incubation of GAR1 with desalted torula yeast RNA (Sigma-Aldrich), NaCl and 1,6-hexanediol, at the indicated concentrations, using the experiment buffer mentioned above. For NaCl and 1,6-hexanediol experiments, 100 μM of GAR1 was used. Data were collected in triplicates and is represented as the mean ± standard deviation (SD).

### Confocal microscopy

Images of *in vitro* GAR1:RNA condensates were acquired using a TCS SP5 inverted confocal microscope (Leica Microsystems CMS) equipped with Plan-Apochromat CS 63x, with NA of 1.2, water immersion objective (Carl Zeiss Microscopy), at room temperature. For the experiments, 99.5 µM of unlabeled GAR1 were mixed with 0.5 µM of GAR1-EGFP in the experiment buffer. Samples of 30 µL were placed on µ-Slide 18 well-flat uncoated microplates (Enzyfarma). GAR1-EGFP samples were excited at 488 nm, and emission was collected between 495 and 600 nm. To monitor RNA co-localization within the droplets, torula yeast RNA (Sigma-Aldrich) was stained with 1 µM of SYTO Deep Red Nucleic Acid Stain (Invitrogen). Images were acquired by simultaneously exciting GAR1-EGFP and RNA-SYTO. The SYTO stain was excited at 633 nm and emission was recorded between 645 and 800 nm. Imaging of SMN TD partitioning was achieved by adding Alexa Fluor 647 SMN TD (99% unlabeled and 1% labeled) to GAR1:RNA condensates. Alexa Fluor 647 was excited at 633 nm and emission was collected between 645 and 800 nm.

Images were collected in triplicate with a 2048 x 2048 resolution, processed and analyzed using ImageJ (Fiji) ^63^.

### FRAP *in vitro* and in live cells

Photobleaching was performed with a 488 nm argon laser. For FRAP experiments of GAR1 *in vitro* condensates, circular regions of interest (ROI) were bleached at the center of droplets that contained a diameter of >5 µm. In live cell nuclei, circular ROI with diameter of 1.9 µm were bleached. For fluorescence correction, additional ROIs were measured under the same conditions as the *in vitro* FRAP experiments. For background correction, fluorescence from a condensate-free area was measured, and to correct for photofading fluorescence was measured in a non-bleached condensate. For FRAP experiments in live cells, fluorescence from outside the cell was used for background correction, and fluorescence across the entire image was used for photofading correction. The fluorescence intensities from bleached ROI were then corrected for background and photofading as previously described ^64^. Normalized recovery curves were fitted to a single exponential FRAP recovery equation, y=A*(1-e^k*t^), where *A* represents the plateau corresponding to the mobile fraction, *k* is the recovery rate and *t* is the recovery time ^64^. The apparent diffusion coefficient and the mobile fractions were estimated as previously described ^64,65^. Microscopy images and FRAP data were collected and processed in ImageJ (Fiji) ^63^. The *in vitro* and *in cell* FRAP fitting were performed using GraphPad Prism 9.3.0 (GraphPad Software Inc.), and R 4.4.1 in RStudio 2024.09.0, respectively.

### NMR spectroscopy

NMR experiments were acquired in a Bruker Avance III and Bruker Avance NEO spectrometers, operating at 600 MHz and 800 MHz ^1^H Larmor frequencies, respectively. Both spectrometers were equipped with Z-gradient cryogenic probes. ^1^H chemical shifts were calibrated through internal referencing using 50 μM DSS (Eurisotop), while ^15^N and ^13^C chemical shifts were referenced indirectly to ^1^H using conversion factors derived from ratios of NMR frequencies ^66^. NMR data were processed using Bruker TopSpin 4.0.6 software (Bruker BioSpin) and analyzed with CARA and SPARKY software ^67,68^. All NMR experiments were performed with samples in the experiment buffer with 10% D_2_O.

To validate the assignment of the SMN^WT^ TD under our experimental conditions, we transferred the previously deposited assignment by Selenko *et al*. ^14^ (BMRB: 4899) and acquired additional 2D and 3D experiments. These included 2D ^1^H-^15^N HSQC, 2D ^1^H-^1^H TOCSY, 2D ^1^H-^1^H NOESY, and 3D HNCO (^15^N-labeled, ^13^C natural abundance). Experiments were performed at 15°C with 430 µM or 250 µM ^15^N-labeled SMN^WT^ TD and SMN^E134K^ TD, respectively. TOCSY and NOESY experiments were acquired with 60 ms and 120 ms mixing times, respectively. Heteronuclear ^15^N {^1^H} NOE experiments were acquired with a relaxation delay of 10 s and analyzed as previously described ^69^.

To probe the interaction between SMN TD and GAR1, we performed titration experiments. Briefly, ^15^N-labeled SMN Tudor WT or E134K variant (fixed 100 µM concentration) were titrated with unlabeled GAR1, and the SMN Tudor ^1^H-^15^N resonance chemical shifts and intensity variations were monitored by recording 2D ^1^H-^15^N HSQC spectra at 15°C. The combined chemical shift perturbations and the chemical shift threshold values were calculated using the equations reported by Schumann *et al.* ^70^. SMN TD structure corresponding to the construct used in this study (UniProt code: Q16637, fragment 83-147) was predicted by AlphaFold 2 ^71^. Model residue-specific electrostatic potential was calculated and displayed with surface coloring using the Coulombic electrostatic potential mode on Chimera X ^72^.

For the RNA competition experiments, samples of 10 µM of ^15^N-labeled SMN^WT^ TD and unlabeled GAR1 (in an equimolar ratio) were progressively titrated with desalted torula yeast RNA (Sigma-Aldrich), in a 1:0.1, 1:0.5 and 1:1 GAR1:RNA mass ratio. The variations in SMN^WT^ TD chemical shifts were monitored by acquiring 2D ^1^H-^15^N HSQC spectra at each RNA concentration, at 25°C.

## Acknowledgements

This work was supported by Fundação para a Ciência e a Tecnologia (FCT-Portugal) for funding UCIBIO project (UIDP/04378/2020 and UIDB/04378/2020) and iBB (UIDB/04565/2020), and Associate Laboratory Institute for Health and Bioeconomy – i4HB project (LA/P/0140/2020). S.S.F. acknowledges FCT-Portugal for the PhD fellowship (PD/BD/148028/2019) under the PTNMRPhD Program. P.O. acknowledges FCT-Portugal for the PhD fellowship (2021.05564.BD). N.A.S.O. acknowledges the support of a fellowship from the ”la Caixa” Foundation (ID 100010434), with the fellowship code LCF/BQ/DR23/12000018. T.S.B. acknowledges the Discovery Fellowship BB/W009242/1 from the Biotechnology and Biological Sciences Research Council (BBSRC). J.Oroz is a recipient of a Leonardo Grant from the Spanish BBVA Foundation (BBM_TRA_0203) and a Ramón y Cajal Grant (RYC2018-026042-I funded by MCIN/AEI/10.13039/501100011033 and by “ESF Investing in your future”). J.Oroz was supported by the Grant PID-2019-109276RA-I00 and D.V.L by the PID-2019-109306RB-I00 and PID2022-137806OB-I00, funded by the Spanish Ministry of Science, Innovation and Universities: MICIN/AEI/10.13039/501100011033/FEDER, UE. The Spanish funding agency had no role in the study design or the data interpretation.

One of the fluorescence microscopy facilities used in this work is integrated in the national infrastructure of PPBI—the Portuguese Platform of Bioimaging.

The NMR spectrometers in UCIBIO are a node of the National NMR Network and are supported by FCT (ROTEIRO/0031/2013 and PINFRA/22161/2016) cofounded by FEDER through COMPETE 2020, POCI, PORL and FCT through PIDDAC. The 800 MHz spectrometer present in the “Manuel Rico” NMR laboratory (LMR-CSIC) is a node of the Spanish Large-Scale National Facility (ICTS R-LRB-MR).

## Author contributions

S.S.F., J.M.A.O, J.O., D.V.L. and E.J.C. conceptualized the project. S.S.F. developed the protocols for recombinant protein production. S.S.F. and P.O. performed the biochemical assays. S.S.F. and F.F. performed the *in vitro* imaging and FRAP experiments. S.S.F., P.O. and D.V.L. performed the NMR experiments. N.A.S.O. and B.R.P. performed cell assays and imaging. N.A.S.O. and T.S.B. performed the FLIM-FRET experiments. Project administration and supervision done by J.M.A.O., J.O., D.V.L., and E.J.C. S.S.F. wrote the original draft. Manuscript review and editing were done by N.A.S.O., J.M.A.O., J.O., D.V.L. and E.J.C. Funding acquired by. J.M.A.O., J.O., D.V.L, and E.J.C. All authors read and approved the manuscript.

