## Supplementary Information for "Pathological mutation in SMN impairs modulation of GAR1 phase separation linking condensate dysfunction to Spinal Muscular Atrophy"

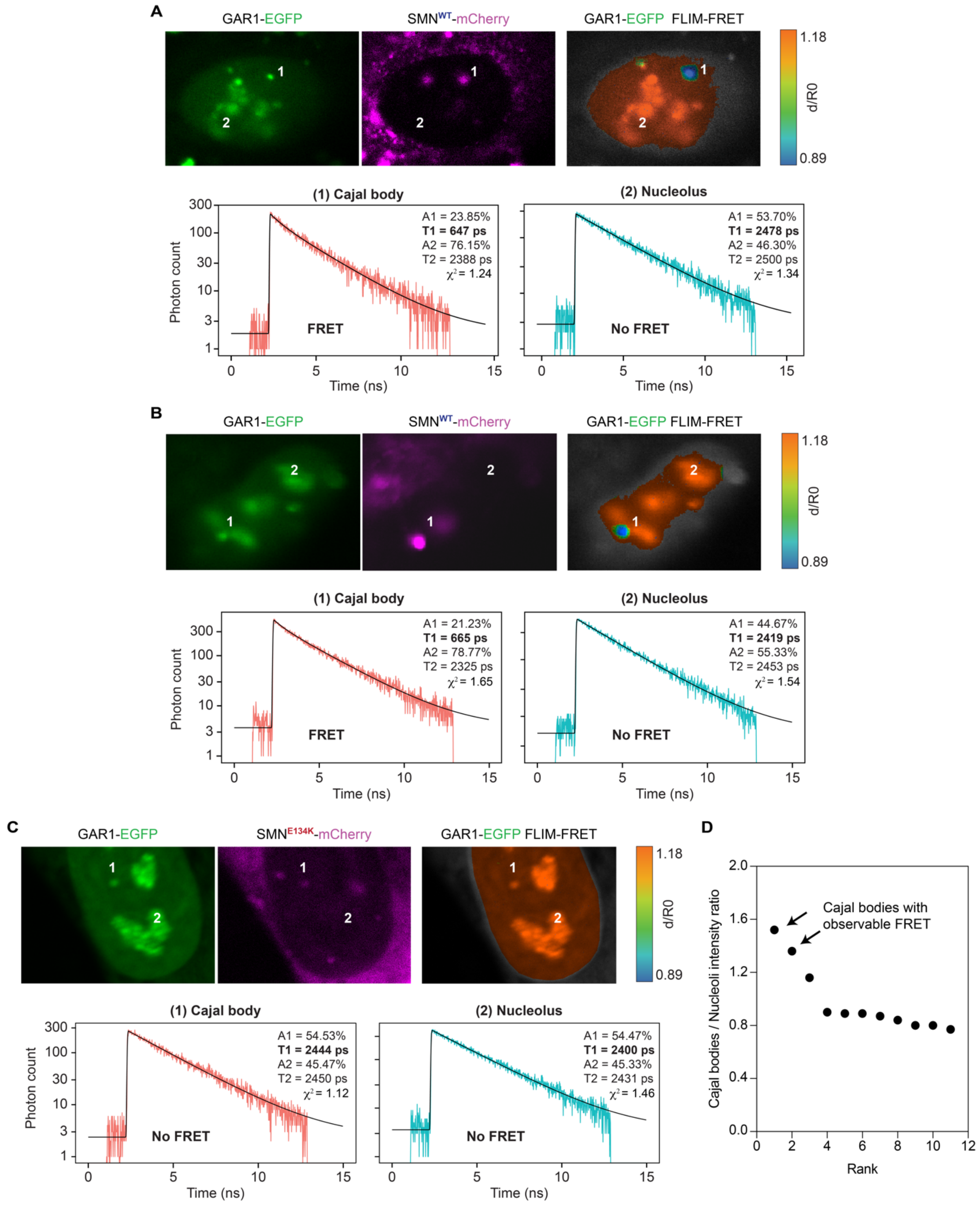


**Figure S1 – FLIM-FRET data supports interaction between GAR1 and SMN$^{WT}$ in Cajal bodies –** Förster resonance energy transfer (FRET) was assessed by fluorescence lifetime imaging (FLIM) of HeLa cells co-transfected with plasmid DNA encoding GAR1-EGFP and mCherry-SMN (WT or E134K mutant). **A,B.** Two cells where FRET can be observed in a Cajal body between GAR1 and SMN$^{WT}$. Representative images of GAR1-EGFP intensity, mCherry-SMN intensity, and lifetime-based donor acceptor distance (d/R0, distance relative to Förster radius) (upper panels). Representative

FLIM curves and double-exponential fit parameters from Cajal bodies or nucleoli (lower panels). **C.** Example cell where no FRET can be observed between GAR1 and $SMN^{E134K}$. Representative images of GAR1-EGFP intensity, mCherry-SMN intensity, and lifetime-based donor acceptor distance (d/R0, distance relative to Förster radius) (upper panel). Representative FLIM curves and double-exponential fit parameters from Cajal bodies or nucleoli (lower panel). **D.** Measured GAR1-EGFP fluorescence intensity in Cajal bodies relative to nucleoli from the same cell, in descending order. We observed FRET in the two Cajal bodies with the highest relative brightness (out of 11), suggesting that signal-to-noise ratio might limit FRET detection [1].

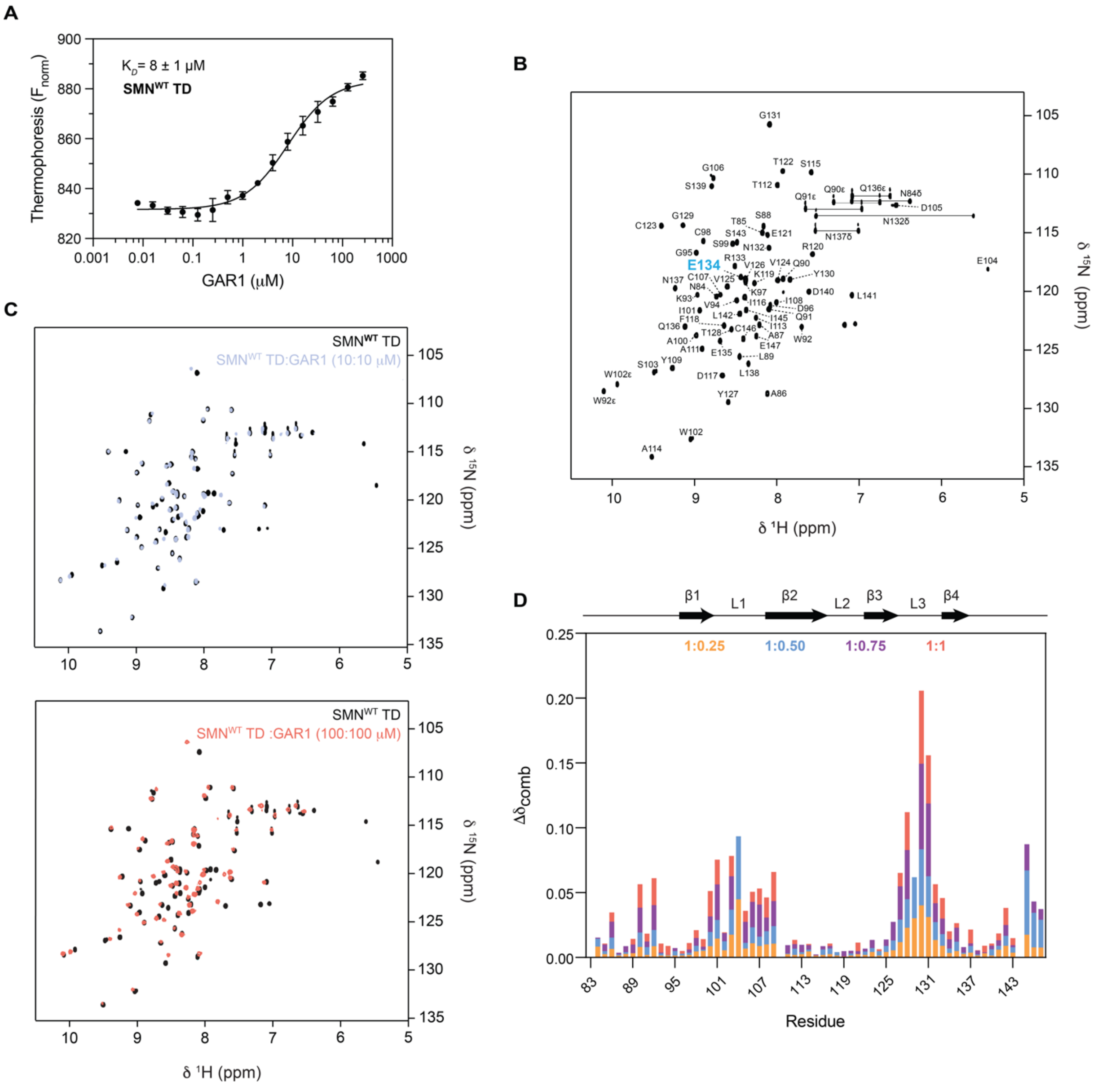

**Figure S2 – MST affinity data and NMR characterization of SMN Tudor domain binding to GAR1 – A.** GAR1 and $SMN^{WT}$ TD binding affinity measured via microscale thermophoresis. Data reported as mean ± SD, *n* = 3 independent experiments. **B.** Resonance assignment of $SMN^{WT}$ TD construct used in our study (UniProt code: Q16637; residues 83-147) validated from deposited data (BMRB: 4899). **C.** Overlay of $^{1}H$-$^{15}N$ HSQC spectra of $^{15}N$-labeled $SMN^{WT}$ TD at 10 μM (upper panel) and 100 μM (lower panel) with unlabeled GAR1 in equimolar ratio. **D.** Overlay of combined chemical shifts of $^{15}N$-labeled $SMN^{WT}$ TD resonances from the titration with unlabeled GAR1, presented as color-coded ratios of $SMN^{WT}$ TD:GAR1. Above the plots is a depiction of the secondary structure of $SMN^{WT}$ TD. β – beta-sheet, L – loop.

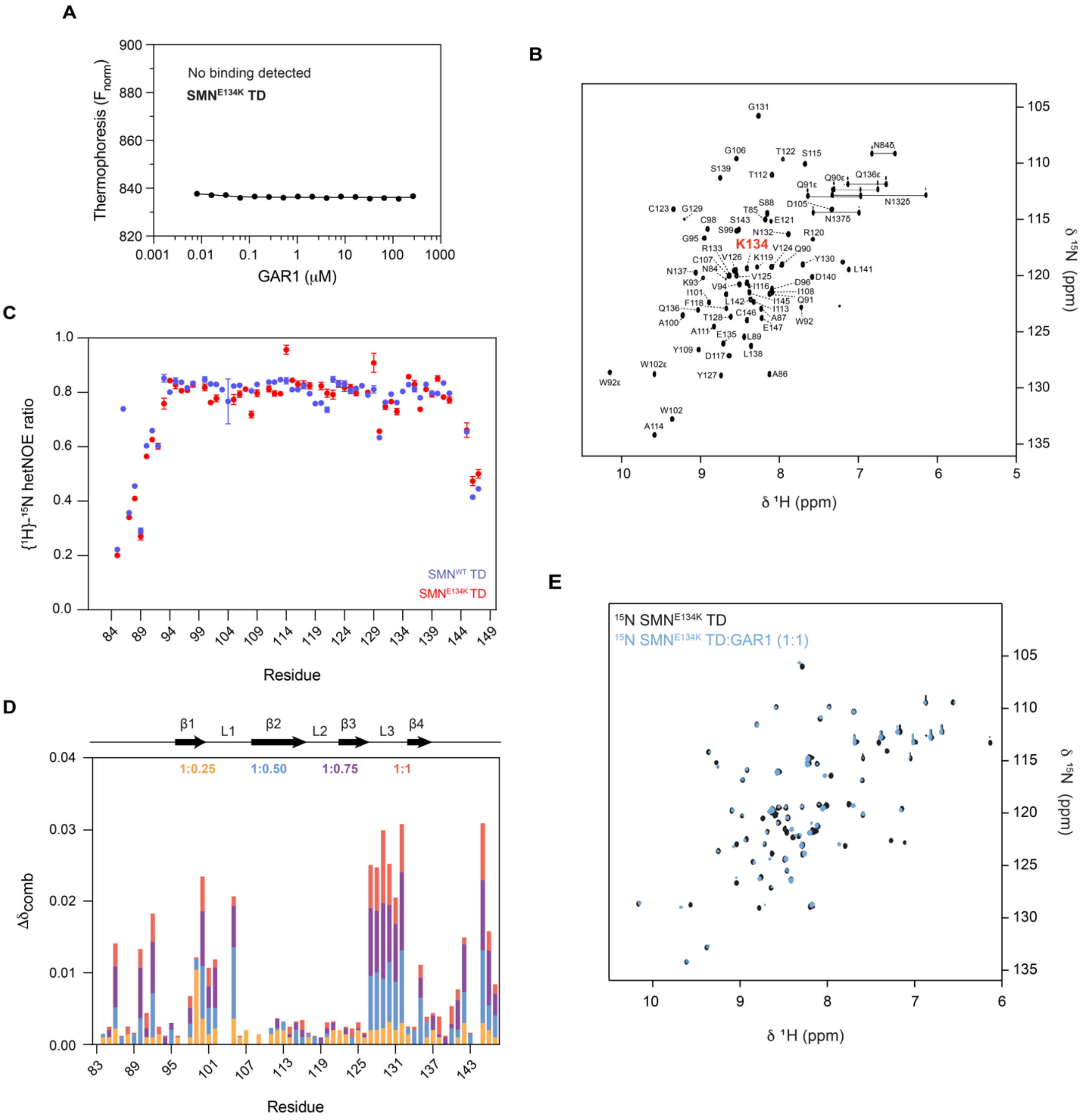


**Figure S3 – MST affinity data and NMR characterization of SMN E134K Tudor domain binding to GAR1 – A.** GAR1 and $SMN^{E134K}$ TD microscale thermophoresis binding assay showing a significant decrease in affinity compared to $SMN^{WT}$ TD. Data reported as mean ± SD, *n* = 3 independent experiments. **B.** Resonance assignment of $SMN^{E134K}$ TD construct used in our study (UniProt code: Q16637; residues 83-147, mutated lysine 134) based on deposited data (BMRB: 4899). **C.** $\{^{1}H\}$-$^{15}N$ hetNOE plot of SMN TD WT and E134K. Error bars, SD calculated from the background signal. **D.** Overlay of combined chemical shifts of $^{15}N$-labeled $SMN^{E134K}$ TD resonances from the titration with unlabeled GAR1, presented as color coded-ratios of $SMN^{E134K}$ TD:GAR1. Above the plots is a depiction of the secondary structure of $SMN^{E134K}$ TD. **E.** Overlay of $^{1}H$-$^{15}N$ HSQC spectra of $^{15}N$-labeled $SMN^{E134K}$ TD without (black) and with unlabeled GAR1 in equimolar ratio (blue). β – beta-sheet, L – loop.

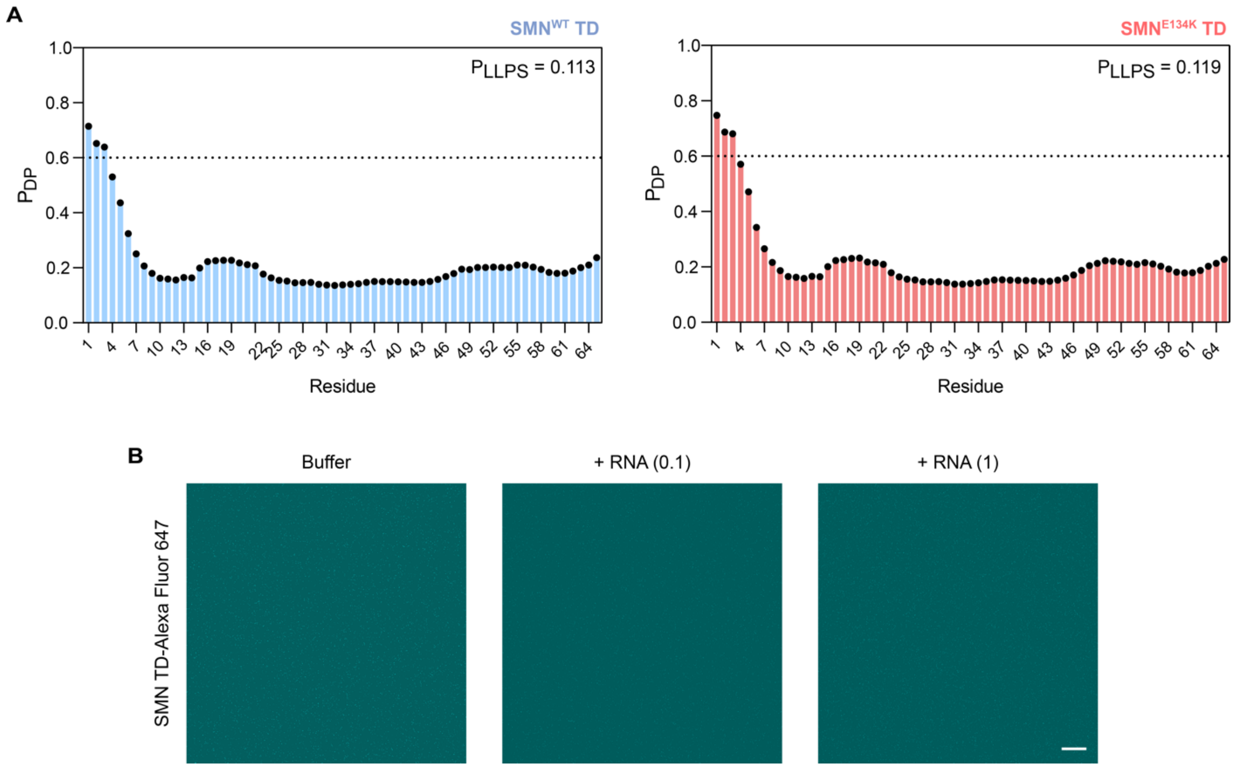


**Figure S4 – SMN Tudor domain does not phase separate *in vitro* - A.** The FuzDrop algorithm predicts that both SMN TD WT (left panel) and E134K variant (right panel) do not have propensity to undergo LLPS, as indicated by the FuzDrop probability scores below the threshold of 0.6. **B.** Confocal fluorescence microscopy control experiments of 100 μM SMN$^{WT}$ TD-Alexa Fluor 647, both in buffer and in the presence of RNA in a 1:0.1 and 1:1 protein:RNA ratio, reveals that SMN TD does not phase separate *in vitro* under our experimental conditions. Scale bar, 10 μm; $P_{LLPS}$ – probability score for spontaneous LLPS; $P_{DP}$ – residue-specific probability for droplet formation.

**Table S1 – Recombinant protein sequences used in this study**

| Construct name | Amino acid sequence |
|---|---|
| $His_6$-MBP-TEV | MGSDKIHHHHHHSSGTKIEEGKLVIWINGDKGYNGLAEVGKKFEKDTGIKVTVEHPDKLEEKFPQVAATGDGPDIIFWAHDRFGGYAQSGLLAEITPDKAFQDKLYPFTWDAVRYNGKLIAYPIAVEALSLIYNKDLLPNPPKTWEEIPALDKELKAKGKSALMFNLQEPYFTWPLIAADGGYAFKYENGKYDIKDVGVDNAGAKAGLTFLVDLIKNKHMNADTDYSIAEAAFNKGETAMTINGPWAWSNIDTSKVNYGVTVLPTFKGQPSKPFVGVLSAGINAASPNKELAKEFLENYLLTDEGLEAVNKDKPLGAVALKSYEEELAKDPRIAATMENAQKGEIMPNIPQMSAFWYAVRTAVINAASGRQTVDEALKDAQTNSGSDITSLYKKAEGGTENLYFQGH |
| GAR1 | MSFRGGGRGGFNRGGGGGGFNRGGSSNHFRGGGGGGGGGNFRGGGRGGFGRGGGRGGFNKGQDQGPPERVVLLGEFLHPCEDDIVCKCTTDENKVPYFNAPVYLENKEQIGKVDEIFGQLRDFYFSVKLSENMKASSFKKLQKFYIDPYKLLPLQRFLPRPPGEKGPPRGGGRGGRGGGRGGGGRGGGRGGGFRGGRGGGGGGFRGGRGGGFRGRGH |
| GAR1-Linker-EGFP | MSFRGGGRGGFNRGGGGGGFNRGGSSNHFRGGGGGGGGGNFRGGGRGGFGRGGGRGGFNKGQDQGPPERVVLLGEFLHPCEDDIVCKCTTDENKVPYFNAPVYLENKEQIGKVDEIFGQLRDFYFSVKLSENMKASSFKKLQKFYIDPYKLLPLQRFLPRPPGEKGPPRGGGRGGRGGGRGGGGRGGGRGGGFRGGRGGGGGGFRGGRGGGFRGRGHGAPGSAGSAAGSGGAPGSAGSAAGSGVSKGEELFTGVVPILVELDGDVNGHKFSVSGEGEGDATYGKLTLKFICTTGKLPVPWPTLVTTLTYGVQCFARYPDHMKQHDFFKSAMPEGYVQERTIFFKDDGNYKTRAEVKFEGDTLVNRIELKGIDFKEDGNILGHKLEYNYNSHKVYITADKQKNGIKVNFKTRHNIEDGSVQLADHYQQNTPIGDGPVLLPDNHYLSTQSALSKDPNEKRDHMVLLEFVTAAGITLGMDELYK |
| $SMN^{WT}$ TD | KNTAASLQQWKVGDKCSAIWSEDGCIYPATIASIDFKRETCVVVYTGYGNR**E**EQNLSDLLSPICE |
| $SMN^{E134K}$ TD | KNTAASLQQWKVGDKCSAIWSEDGCIYPATIASIDFKRETCVVVYTGYGNR**K**EQNLSDLLSPICE |

**Table S2 – Protein sequences for transient transfection in cells**

| Construct name | Amino acid sequence |
|---|---|
| GAR1-Linker-EGFP | MSFRGGGRGGFNRGGGGGGFNRGGSSNHFRGGGGGGGGGNFRGGGRGGFGRGGGRGGFN<br>KGQDQGPPERVVLLGEFLHPCEDDIVCKCTTDENKVPYFNAPVYLENKEQIGKVDEIFGQLRDFYFS<br>VKLSENMKASSFKKLQKFYIDPYKLLPLQRFLPRPPGEKGPPRGGGRGGRGGGRGGGGRGGGRG<br>GGFRGGRGGGGGGFRGGRGGGFRGRGHGAPGSAGSAAGSGGAPGSAGSAAGSGVSKGEELFT<br>GVVPILVELDGDVNGHKFSVSGEGEGDATYGKLTLKFICTTGKLPVPWPTLVTTLTYGVQCFARYPD<br>HMKQHDFFKSAMPEGYVQERTIFFKDDGNYKTRAEVKFEGDTLVNRIELKGIDFKEDGNILGHKLEY<br>NYNSHKVYITADKQKNGIKVNFKTRHNIEDGSVQLADHYQQNTPIGDGPVLLPDNHYLSTQSALSK<br>DPNEKRDHMVLLEFVTAAGITLGMDELYK |
| mCherry-Linker-SMN$^{WT}$ | MVSKGEEDNMAIIKEFMRFKVHMEGSVNGHEFEIEGEGEGRPYEGTQTAKLKVTKGGPLPFAWDIL<br>SPQFMYGSKAYVKHPADIPDYLKLSFPEGFKWERVMNFEDGGVVTVTQDSSLQDGEFIYKVKLRGT<br>NFPSDGPVMQKKTMGWEASSERMYPEDGALKGEIKQRLKLKDGGHYDAEVKTTYKAKKPVQLPGA<br>YNVNIKLDITSHNEDYTIVEQYERAEGRHSTGGMDELYKGAPGSAGSAAGSGMAMSSGGSGGGVP<br>EQEDSVLFRRGTGQSDDSDIWDDTALIKAYDKAVASFKHALKNGDICETSGKPKTTPKRKPAKKNKS<br>QKKNTAASLQQWKVGDKCSAIWSEDGCIYPATIASIDFKRETCVVVYTGYGNREEQNLSDLLSPICE<br>VANNIEQNAQENENESQVSTDESENSRSPGNKSDNIKPKSAPWNSFLPPPPPMPGPRLGPGKPGL<br>KFNGPPPPPPPPPPPHLLSCWLPPFPSGPPIIPPPPPICPDSLDDADALGSMLISWYMSGYHTGYYM<br>GFRQNQKEGRCSHSLN |
| mCherry-Linker-SMN$^{E134K}$ | MVSKGEEDNMAIIKEFMRFKVHMEGSVNGHEFEIEGEGEGRPYEGTQTAKLKVTKGGPLPFAWDIL<br>SPQFMYGSKAYVKHPADIPDYLKLSFPEGFKWERVMNFEDGGVVTVTQDSSLQDGEFIYKVKLRGT<br>NFPSDGPVMQKKTMGWEASSERMYPEDGALKGEIKQRLKLKDGGHYDAEVKTTYKAKKPVQLPGA<br>YNVNIKLDITSHNEDYTIVEQYERAEGRHSTGGMDELYKGAPGSAGSAAGSGMAMSSGGSGGGVP<br>EQEDSVLFRRGTGQSDDSDIWDDTALIKAYDKAVASFKHALKNGDICETSGKPKTTPKRKPAKKNKS<br>QKKNTAASLQQWKVGDKCSAIWSEDGCIYPATIASIDFKRETCVVVYTGYGNRKEQNLSDLLSPICE<br>VANNIEQNAQENENESQVSTDESENSRSPGNKSDNIKPKSAPWNSFLPPPPPMPGPRLGPGKPGL<br>KFNGPPPPPPPPPPPHLLSCWLPPFPSGPPIIPPPPPICPDSLDDADALGSMLISWYMSGYHTGYYM<br>GFRQNQKEGRCSHSLN |
